# Modeling the complete kinetics of coxsackievirus B3 reveals human determinants of host-cell feedback

**DOI:** 10.1101/2020.07.26.222174

**Authors:** Aaron B. Lopacinski, Andrew J. Sweatt, Christian M. Smolko, Elise Gray-Gaillard, Cheryl A. Borgman, Millie Shah, Kevin A. Janes

## Abstract

Complete kinetic models are pervasive in chemistry but lacking in biological systems. We encoded the complete kinetics of infection for coxsackievirus B3 (CVB3), a compact and fast-acting RNA virus. The kinetics are built from detailed modules for viral binding–delivery, translation–replication, and encapsidation. Specific module activities are dampened by the type I interferon response to viral double-stranded RNAs (dsRNAs), which is itself disrupted by viral proteinases. The validated kinetics uncovered that cleavability of the dsRNA transducer mitochondrial antiviral signaling protein (MAVS) becomes a stronger determinant of viral outcomes when cells receive supplemental interferon after infection. Cleavability is naturally altered in humans by a common MAVS polymorphism, which removes a proteinase-targeted site but paradoxically elevates CVB3 infectivity. These observations are reconciled with a simple nonlinear model of MAVS regulation. Modeling complete kinetics is an attainable goal for small, rapidly infecting viruses and perhaps viral pathogens more broadly.

## INTRODUCTION

In chemistry, a complete kinetic model specifies products and reactants, their stoichiometry, any catalysts, and the step-by-step sequence of key elementary reactions from start to finish (Moore et al., 1981). Direct analogies in biological systems are hard to identify because the steps are uncertain and the definition of “start” or “finish” is often arbitrary. Not so for picornaviruses, a family of non-enveloped, positive-strand RNA viruses that infect, rapidly amplify their genetic material and capsid proteins, and lyse the host cell marking termination. Steps in the picornaviral life cycle are fully delineated (Racaniello, 2013). The translation of the coding RNA genome as a single polypeptide ensures an equal proportion of each viral protein during synthesis. From the standpoint of products and reactants, picornaviruses present an opportunity to construct complete kinetic models.

The power of kinetic mechanisms lies in the explanatory and predictive models that they generate (Moore et al., 1981). Mathematical models of viral translation–replication have a rich history, but they are formally incomplete in omitting the details of early binding–entry, intermediate antiviral pathways, and late self-assembly of viral particles (Yin and Redovich, 2018). Some omissions are for simplicity and scope, but others are for lack of biological parameters or quantities. As human disease agents, picornaviruses within the *Enterovirus* genus—which includes rhinovirus, poliovirus, and coxsackievirus strains—have been parameterized extensively (Racaniello, 2013). Enteroviruses thus represent the best-case testbed for proposing a complete kinetic model of a biological process if one can be defined.

Here, we drafted the complete kinetics for coxsackievirus B3 (CVB3), an enterovirus that infects the heart and drives viral myocarditis (Esfandiarei and McManus, 2008). With attention to molecular detail, we encoded interacting modules for viral delivery, translation–replication, and encapsidation, overlaying a prototypical antiviral response along with viral antagonism of that response (Figure 1A). Each component presented its own challenges, which we tackled module-by-module before simulating the complete kinetics of CVB3 infection. In a cardiomyocyte-derived cell line, the integrated modules captured host-cell susceptibility and viral RNA–protein dynamics with little-to-no parameter fitting of specific mechanisms. The generalized antiviral and antagonistic feedbacks were screened combinatorially to identify time-dependent interactions later verified experimentally. Host and viral feedbacks converged on an important role for MAVS, a polymorphic host-cell transducer of the antiviral response to dsRNA that is cleaved by an enterovirus-encoded proteinase (Mukherjee et al., 2011). The centrality of MAVS led to the discovery of a common variant in humans (MAVS Q93E), which reroutes cleavage during CVB3 infection and favors viral propagation. The net effect of this polymorphism on feedback arises from nonlinearities in MAVS regulation, as illustrated by a standalone model of its provisional activation mechanism. For virology, complete kinetics provide a systems-level end goal for decades of research and possibly an aid to prioritizing scientific activity during viral outbreaks.

**Figure 1.**
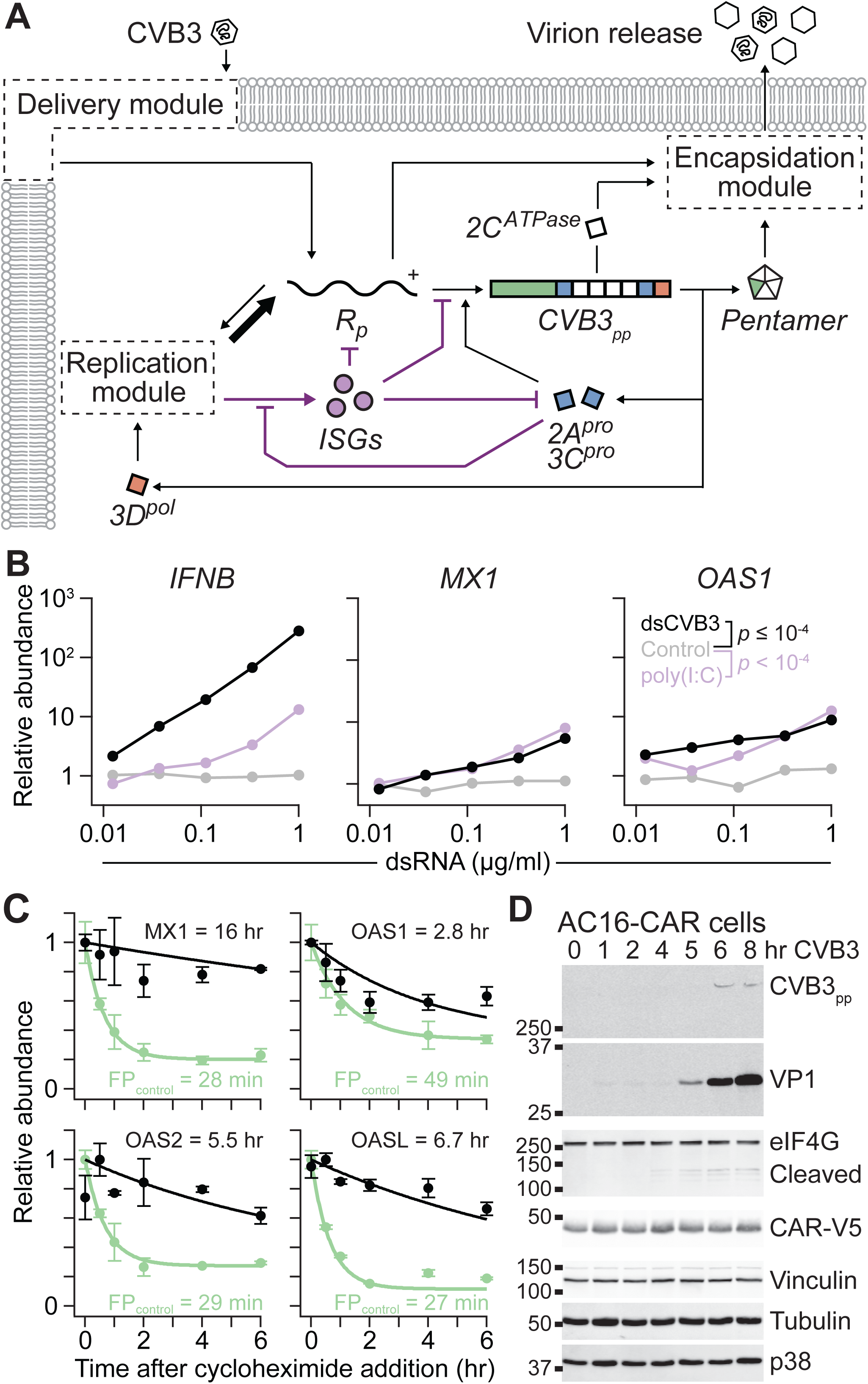
A modular encoding of the CVB3 life cycle elaborated with antiviral responses and viral antagonism of host-cell processes. (A) Overview of the CVB3 model architecture. CVB3 enters through a module of receptors and trafficking states that delivers its positive-strand RNA genome (*R_p_*) to the cytoplasm. *R_p_* is translated into a polyprotein (*CVB3_PP_*) that matures into capsid subunits contributing one fifth of a subunit (*Pentamer*) in a 12-subunit assembly, enteroviral proteinases (*2A^pro^*–*3C^pro^*), hydrophobic proteins (*2C^ATPase^*), and RNA-dependent RNA polymerase (*3D^pol^*). *3D^pol^* replicates *R_p_* through a negative-strand intermediate in a module that gives rise to excess *R_p_*, which joins with *Pentamer* in an encapsidation module that self-assembles *Pentamer* around *R_p_* and leads to virion release. Interferon stimulated genes (*ISGs*) are induced as a consequence of the dsRNA associated with CVB3 replication and impede viral progression where indicated; viral sensing is also antagonized by *3C^pro^* (Mukherjee et al., 2011). Expanded descriptions of the delivery, replication, and encapsidation modules are shown in Figures 2A, 3A, 4A, and 4D, and model parameters are available in Table S1. (B) Transfected dsCVB3 elicits a robust type I interferon response. AC16-CAR cells were lipofected with *n* = 5 doses of the indicated dsRNAs for four hours and analyzed for the indicated ISGs with *PRDX6*, *HINT1*, and *GUSB* used for loading normalization. Differences between conditions were assessed by Šidák-corrected, log-transformed three-way ANOVA. Uninduced target genes (*EIF2AK2* and *GAPDH*) are analyzed in Figure S1A. (C) ISGs are long lived when translated. 293T/17 cells were lipofected with V5 epitope-tagged plasmids, treated with 50 µM cycloheximide for the indicated times, and lysed for quantitative immunoblotting. Half-lives for the ISGs MX1, OAS1, OAS2, OASL, and FP_control_ (a fast-degrading protein fragment used as a positive control) were estimated by nonlinear least-squares curve fitting. Data are shown as the mean ± range of biological duplicates at *n* = 6 different time points. Representative immunoblots are shown in Figure S1B. (D) CVB3 polyprotein maturation coincides with cleavage of host-cell targets. AC16-CAR cells were infected with CVB3 at MOI = 10 for the indicated times and immunoblotted for VP1 capsid protein (also present in the full-length polyprotein [CVB3_pp_]) and eIF4G [a host protein cleaved by *2A^pro^* (Haghighat et al., 1996)] with ectopic CAR-V5, vinculin, tubulin, and p38 used as loading controls. The image gamma was adjusted for polyprotein–VP1 (gamma = 20) and eIF4G (gamma = 2) to show band pairs at the same exposure. See also Figure S1 and Table S1.

## RESULTS

### Modular draft of a complete kinetic model for the CVB3 life cycle

We pursued a complete kinetic model for acute CVB3 infection by deconstructing its life cycle into separable modules that could be developed independently (Figure 1A). The cellular tropism of CVB3 is determined by specific cell-surface receptors (Bergelson et al., 1997; Shafren et al., 1995), which internalize the virus before endosomal escape of its positive-strand RNA genome (*R_p_*) into the cytoplasm. From here, the intracellular steps generalize to all enteroviruses: i) translation of *R_p_* into polyprotein (*CVB3_PP_*), ii) maturation of *CVB3_PP_* into capsid subunits and nonstructural proteins for the virus, iii) cis replication of *R_p_* through a negative-strand intermediate by RNA-dependent RNA polymerase (*3D^pol^*) localized to host-cell membranes hijacked by the virus, and iv) encapsidation of *R_p_* around 12 capsid pentamers aided by hydrophobic proteins (*2C^ATPase^*) and concluding with lytic release from the host (Baggen et al., 2018). In contrast to cell-signaling pathways (Friedman and Perrimon, 2007), the viral conduits to and from these modules are clearly defined, and an acute infection usually completes within hours if unimpeded.

At the nexus of the enteroviral life cycle is the type I interferon response. Viral replication is associated with long, double-stranded replicative intermediates of CVB3 (dsCVB3) that are sensed by innate antiviral pathways like other dsRNAs [e.g., poly(I:C)] (Wang et al., 2010). Cytosolic dsCVB3 gives rise to induction of *IFNB*, autocrine–paracrine signaling, and the induction of interferon-stimulated genes (ISGs) (Feng et al., 2012), such as oligoadenylate synthetases and Mx-family GTPases (Figure 1B). There are hundreds of different ISGs, several of which inhibit core enteroviral processes at multiple points (Figure 1A) (Schoggins et al., 2011). ISGs are predicted to persist for the duration of an infection (Figure 1C), and active type I interferon signaling is a potent inhibitor of enteroviral replication in vivo (Deonarain et al., 2004; Liu et al., 2005). The enteroviral proteinases (*2A^pro^–3C^pro^*)—which mature the *CVB3_pp_* polyprotein and steal host-cell ribosomes for cap-independent viral translation—interfere with activation of the interferon response by cleavage of dsRNA sensors and transducers (Figure 1A) (Feng et al., 2014; Haghighat et al., 1996; Mukherjee et al., 2011). Cleavage of host-cell proteins occurs on the same time scale as polyprotein maturation (Figure 1D) (Etchison et al., 1982; Mukherjee et al., 2011), requiring us to overlay a network of ISG-related negative feedbacks on the CVB3 life cycle (Figure 1A, purple).

The viral life-cycle modules and antiviral–antagonistic feedbacks were encoded as a system of 54 differential equations that are derived in the STAR Methods. Experimental evidence for 91% of the 92 parameters is provided in Table S1. In general engineering design, modules are appealing because they enable individual components to be developed and characterized before integration (Janes et al., 2017). Accordingly, our results are communicated to retain the modular organization of the CVB3 model. For individual modules, we expand biochemical mechanisms, describe critical assumptions or considerations, compare with experiments, and conclude with non-obvious computational predictions.

### Explanatory modeling of CVB3 tropism requires careful extracellular bookkeeping

A hurdle to modeling the complete kinetics of CVB3 arises even before the virus has encountered a host cell. Stock preparations of enterovirus contain many more RNA-filled viral particles than infectious plaque-forming-units (PFUs). Encapsidated RNA genomes often do not reach a host-cell ribosome for translation, and some of those that do contain deleterious mutations from the prior replication. These particles are “defective” from the standpoint of infection, yet they are generally able to bind cell-surface receptors and internalize (Brandenburg et al., 2007). Defective particles thus contribute viral protein and RNA to the system, which must be accounted for at the start of the life cycle.

Particle-to-PFU ratios vary widely within an enterovirus species—the ratio for poliovirus, for example, is reported to be 30:1 to 1000:1 (Racaniello, 2013). We quantified sedimentable RNA content from separately purified CVB3 preparations of known PFU titer and estimated a particle-to-PFU ratio of 800 ± 200 (*n* = 4 preparations; STAR Methods). In the model, we assumed an 800-fold excess of defective particles, which traffic identically to PFUs but do not translate upon entering the host-cell cytoplasm (STAR Methods). The assumption creates a kinetic dead-end for defective particles without any additional kinetic parameters (Table S1). More complicated defective-particle fates yielded similar infection outcomes, as shown after the model was fully developed (see below).

A second challenge relates to the front-end model implementation of viral delivery. The early steps of virus binding and internalization are discrete and thus intrinsically stochastic (Ellis and Delbruck, 1939). However, one negative-strand template can yield hundreds of protein-coding positive strands within an hour (Castro et al., 2007), indicating rapid transition to a regime where deterministic modeling is valid. We balanced these tradeoffs by postulating that the dominant stochasticity was the Poisson noise from the number of PFUs encountered at a given multiplicity of infection (MOI) describing the average PFU per cell within a population (Ellis and Delbruck, 1939). Cell-to-cell variation in all downstream processes was modeled by lognormal random variables centered around the best estimates from the literature (Table S1) and sampled with a user-defined coefficient of variation. We configured the simulations to run in one of two modes: 1) a “single-cell” mode, in which a discrete number of PFUs is simulated with lognormally distributed downstream parameters, and 2) a “cell-population” mode, in which a lognormal instance of the life cycle is paired with a PFU integer drawn from a Poisson distribution about a continuous MOI representing the average. In either mode, dynamic trajectories were well summarized by ∼100 separate iterations. The dual implementation thereby models the average-cell response to an initial condition that is highly stochastic.

Binding and delivery of CVB3 requires decay accelerating factor (DAF, officially named CD55) and coxsackievirus and adenovirus receptor (CAR, officially named CXADR) (Bergelson et al., 1997; Shafren et al., 1995). DAF is a low-affinity receptor (K_D_ ∼ 3 µM) whose transcript is widely expressed at moderate abundance in human tissues (median transcripts per million [TPM] ∼ 25) (Consortium, 2015; Lea et al., 1998). CAR is the high-affinity receptor (K_D_ ∼ 0.2 µM) that is less abundant overall (median TPM ∼ 3.5) and restricted to the heart, brain, and epithelial tissues that CVB3 infects (Consortium, 2015; Goodfellow et al., 2005). DAF is a GPI-linked surface protein that diffuses freely in cholesterol-rich microdomains on the cell surface, whereas CAR is a cell-cell adhesion protein localized to tight junctions. Binding of CVB3 to DAF enables the CVB3:DAF complex to traffic to tight junctions, where CVB3 is bound by CAR to promote endosomal internalization (Figure 2A) (Coyne and Bergelson, 2006). In the model, we assumed that CAR is instantly degraded upon CVB3 internalization at characteristic rates for endocytosis (Chung et al., 2005). However, results in permissive hosts were unchanged if CAR was instead recycled instantaneously to the surface (Figure S2A). Rates of rapid endosomal escape were estimated from live-cell experiments with labeled poliovirus (Brandenburg et al., 2007). The output of the delivery module is positive-strand RNA (both infectious and defective) in the cytoplasm.

**Figure 2.**
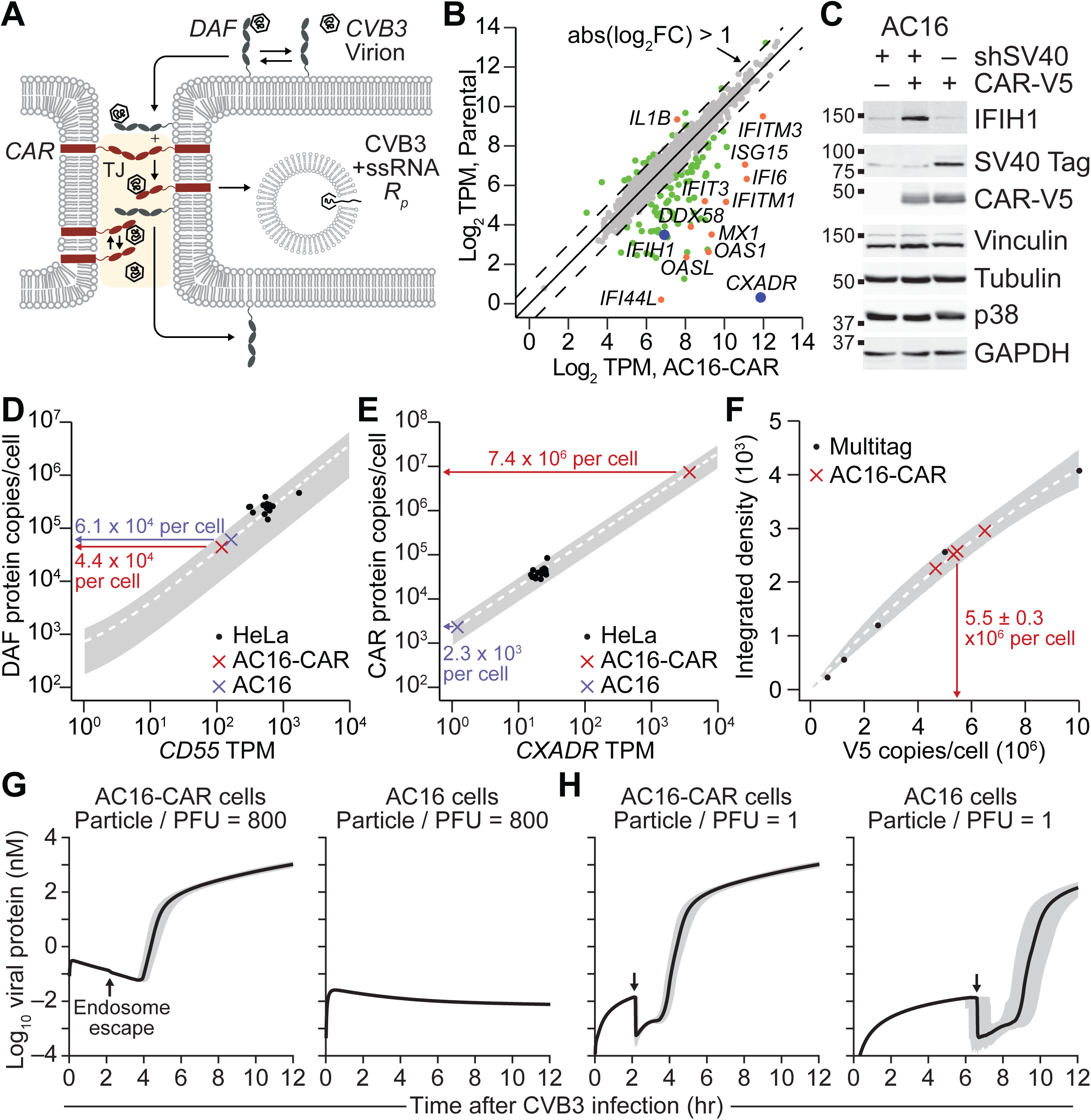
Stoichiometric estimation and simulation of CVB3 particles, cargo, and surface receptors. (A) Overview of the CVB3 delivery module. CVB3 binds to DAF and translocates to tight junctions (TJ, beige). In the tight junction, CVB3 unbinds DAF and binds to CAR by two parallel mechanisms (STAR Methods). CAR-bound CVB3 is internalized and DAF is returned to the plasma membrane. At any point, CVB3 may dissociate from its receptors (reverse arrows). After internalization, the viral genome (*R_p_*) escapes the endosome into the cytoplasm. (B) Ectopic expression of CAR in quiescent AC16 cells causes chronic overexpression of immune response genes. Differentially expressed transcripts between quiescent parental AC16 cells (Parental) and CAR-overexpressing AC16 cells (AC16-CAR) (*q* < 0.05; gray) that also have a log_2_ fold change (FC) greater than ±1 (green) and are immune-regulated genes (orange). Proteins encoded by *IFIH1* and *CXADR* (blue) are independently measured in (C). (C) The elevated interferon response of AC16-CAR cells is restricted to the quiescence protocol involving SV40 knockdown. AC16 cells with or without quiescence (shSV40) or CAR overexpression (CAR-V5) were immunoblotted for IFIH1, SV40 T antigen (Tag), and ectopic CAR-V5 with vinculin, tubulin, p38, and GAPDH used as loading controls. Replicated densitometry is shown in Figures S2B–D. (D and E) Estimating DAF–CAR abundance through coupled transcriptomics and proteomics of HeLa cells (Liu et al., 2019). Data from *n* = 14 HeLa variants was used to determine a hyperbolic-to-linear fit (STAR Methods) that was combined with the AC16 transcriptomic data to estimate the per-cell DAF–CAR protein abundances for AC16 cells (blue) and AC16-CAR cells (red). The best-fit curve (white dashed) is shown ± 99% confidence interval of the fit (gray). (F) Direct measurement of CAR abundance in AC16-CAR cells. Recombinant V5-containing Multitag was used to calibrate a V5 quantitative immunoblot and estimate the total per-cell abundance of CAR-V5. Data are shown as the mean ± s.e.m. of *n* = 4 biological replicates (red). The best-fit calibration curve (white dashed) is shown ± 90% confidence interval of the fit (gray). (G and H) Complete kinetics requires accounting of defective CVB3 particles to capture host-cell tropism. For these simulations, DAF abundance was averaged to 52,500 copies per cell. Predictions are shown as the median simulation ± 90% nonparametric confidence interval from *n* = 100 simulations of single-cell infections at 10 PFU with a parameter coefficient of variation of 5%. In (H), the lack of defective particles exaggerates the drop in viral protein associated with endosomal escape (arrows; STAR Methods). Results from (G) and (H) were unchanged if endogenous CAR was estimated from a calibration including the protein estimates from AC16-CAR cells in (F). See also Figure S2.

As a representative host cell, we used AC16 cells, which were originally immortalized by cell fusion of adult ventricular cardiomyocytes with SV40-transformed fibroblasts (Davidson et al., 2005). AC16 cells are not very permissive to CVB3 infection, but they become highly susceptible upon ectopic expression of CAR (Shah et al., 2017). We originally intended to use CAR-overexpressing AC16 cells (AC16-CAR) under quiescent conditions in which SV40 was knocked down and serum reduced (Davidson et al., 2005). However, RNA sequencing (RNA-seq) revealed that CAR overexpression in quiescent cells caused a considerable upregulation of ISGs (Figure 2B), which would confound the analysis. Follow-up experiments revealed that ISG upregulation was specific to the quiescence protocol (Figures 2C and S2B–D), enabling the use of proliferating AC16-CAR cells for the study.

The delivery module is composed entirely of low copy-number events and intermediate species that are difficult to assess experimentally. Consequently, we selected DAF and CAR abundance for a validity check on the module, using the other modules to amplify the effect of CVB3 receptors on viral outputs that were measurable. To parameterize initial conditions for DAF and CAR, we derived estimates from the AC16 RNA-seq data by using an RNA-to-protein conversion factor for each receptor estimated from a panel of different HeLa lines profiled by transcriptomics and proteomics (Edfors et al., 2016; Liu et al., 2019). HeLa cells are highly susceptible to CVB3 and are widely used for virus propagation; thus, per-cell receptor abundances should be in the operating range for productive infection. Both conversions were adequately described by a hyperbolic-to-linear fit providing approximate protein estimates for parental AC16 and AC16-CAR cells (Figures 2D and 2E; STAR Methods). However, CAR estimates from corresponding *CXADR* mRNA in AC16-CAR cells required an enormous extrapolation of the fit (Figure 2E), prompting an independent measurement. Ectopic CAR contains a V5 tag on the C-terminus, which prevents the construct from transmitting adhesion signals to PDZ domain-containing proteins (Excoffon et al., 2004) and enables absolute epitope quantification. Using a recombinant multitag protein containing a V5 epitope, we calibrated V5 immunoblotting to protein copies and interpolated CAR-V5 copies for known numbers of AC16-CAR cells (Figures 2F and S2E). The ∼5.5 million CAR-V5 molecules per cell falls within the prediction interval for the RNA-to-protein conversion fits, supporting that fit-based estimates for DAF (∼61,000 [AC16] and ∼44,000 [AC16-CAR] per cell) and endogenous CAR (∼2300 per cell) are realistic.

Taking an average DAF abundance between the two lines, we simulated the quantitative importance of CAR at endogenous and ectopic levels with a CVB3 infection of 10 PFU. When viral particles were fully accounted for, the model predicted a robust infection in CAR- overexpressing cells but not parental AC16 cells, reflected by viral protein 1 (VP1) expression within 10 hours (Figures 1D and 2G). Host permissiveness was predicted to occur at ∼7000 CAR copies per cell (Figure S2F), a threshold surpassed by all HeLa variants in the panel (Figure 2E) (Liu et al., 2019). By contrast, when defective particles were ignored and the same host-cell conditions simulated, productive CVB3 infections occurred at both ectopic and endogenous abundances of CAR (Figure 2H). Lack of receptor competition from defective particles enables DAF-bound PFUs to access the small number of CARs, internalize, and replicate. Competition was consequential at particle-to-PFU ratios of 200 or greater (Figure S2G), partially overlapping with the range of ratios documented for enteroviruses (Racaniello, 2013). The results suggest that host-cell permissiveness is conditioned on PFU purity relative to defective interfering particles.

### A two-phase encoding of membrane replication predicts quantifiable positive–negative strands and replicative intermediates

After entering the cytoplasm, an infectious positive-strand RNA genome is translated by host-cell ribosomes. In cell-free systems, enteroviral RNA is recognized by 2–5 ribosomes within 15 seconds (Kempf and Barton, 2008). We conservatively assumed a polysome size of 2–3 ribosomes to account for ribosome competition with host-cell mRNAs early in infection. In the model, polysomes increase the net translation of viral proteins along with the retention of positive strands in translation complexes (STAR Methods). Modeling polysomes was critical for appropriately timing the onset of exponential viral protein synthesis under realistic translation rates (Figure S3A).

Any positive strand released from a translation complex was instantly placed on a 3D^pol^- containing surface to capture the cis coupling of enteroviral translation and replication (Figure 3A; STAR Methods) (Goodfellow et al., 2000; Novak and Kirkegaard, 1994). During infection, host-cell membranes are redirected to create molecular factories for genome replication from a negative-strand template (Cuconati et al., 1998). The formation of such “viral replication organelles” (VROs) is widely documented in positive-strand RNA viruses and may serve multiple functions (Miller and Krijnse-Locker, 2008). Using the model, we considered three functions for VROs: 1) shield viral RNAs from degradation, 2) shield viral replicative intermediates from dsRNA sensing, and 3) accelerate replicative processes by concentrating species on a surface. Recent ultrastructural studies suggest a per-cell VRO surface area of 160–185 µm^2^ (Melia et al., 2019), which roughly agrees with our own brightfield estimates of 120 µm^2^ (STAR Methods). These numbers and the ∼7-nm height of a 3D^pol^ enzyme (PDB ID: 3CDW) yield a >2000-fold concentrating effect of associating cytoplasmic viral molecules in the local volume of a VRO surface. Whereas setting either viral RNA degradation or dsRNA sensing on VROs to zero had a negligible impact on the timing of acute infection, reducing the VRO-concentrating effect to less than 100-fold yielded no net output (Figure 3B). Other VRO functions might be important for RNA viruses that replicate more slowly and organize VROs differently (Binder et al., 2013), but these computational results strongly suggest that the dominant role for enteroviral VROs is to accelerate biochemistry.

**Figure 3.**
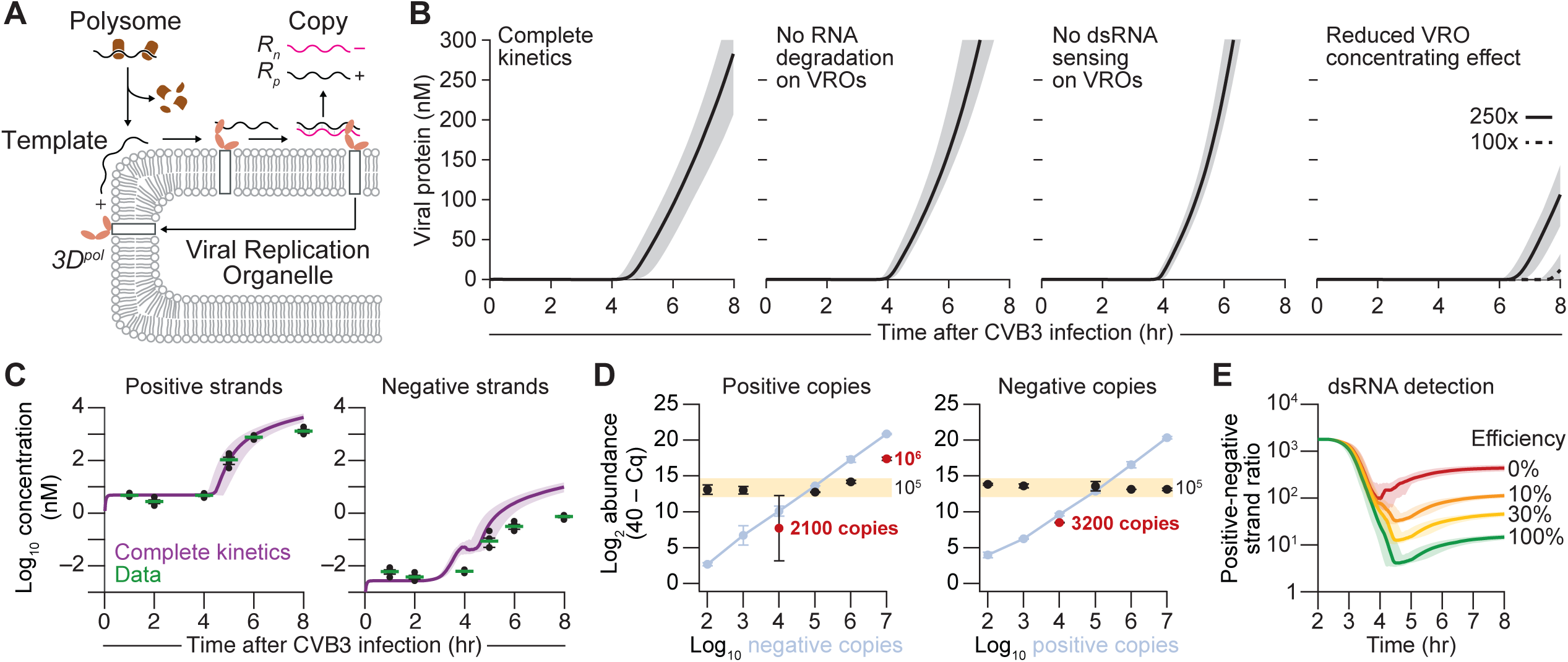
Explosive genome replication requires intracellular membranes and hides negative-strand templates from quantitation. (A) Overview of the replication module. After dissociating from polysomes, the positive-strand RNA genome is released to associate with the surface of viral replication organelles (VROs), where translated RNA-dependent RNA polymerase (*3D^pol^*) resides bound to 3AB (white box). Replication proceeds on the VRO surface, releasing one positive strand (*R_p_*) and one negative strand (*R_n_*). The reciprocal process occurs with *R_n_* as template, except that *R_n_* is not released from *3D^pol^* (STAR Methods). (B) VROs are surface accelerants. Translational output from complete kinetics altered to assume zero RNA degradation, zero dsRNA sensing, or limited concentrating effect on the VRO surface. The concentrating effect in the complete kinetic model is 3216x (see STAR Methods). (C) Complete kinetics captures the absolute viral RNA dynamics of positive and negative strands. Predictions (purple) were compared to data (green) obtained by strand-specific tagged quantitative PCR (qPCR) with purified standards. Data are shown as the geometric mean ± log-transformed standard error of *n* = 4 biological replicates of AC16-CAR cells infected at MOI = 10 for the indicated times. Population-level simulations are shown in Figure S3D. (D) Strand competition between sense and antisense CVB3 genomes in vitro. Strand-specific tagged qPCR of 10^5^ positive copies (left) or negative copies (right) amidst the indicated abundance of complementary strand on the x-axis (measured in blue). Data [black (with outliers highlighted in red) and blue] are shown as the mean log_2_ relative abundance [40 – qPCR quantification cycle (Cq)] (Singh et al., 2019) ± range of assay duplicates at *n* = 6 separate positive-negative strand mixtures. (E) Observed positive-negative strand ratio depends critically on the detection efficiency of dsRNA replicative intermediates. The complete kinetic model was simulated and inventoried with different fractional contributions of dsRNA to the positive- and negative-strand totals. For (B), (C), and (E), predictions are shown as the median simulation ± 90% nonparametric confidence interval from *n* = 100 simulations of single-cell infections at 10 PFU with a parameter coefficient of variation of 5%. See also Figure S3.

Next, we needed to define when the VRO acceleration was triggered kinetically. One 3D^pol^ enzyme by itself is dilute whether in the cytoplasm or on a membrane, but local surface patches of enzyme and positive strand could exhibit acceleration even though the whole-cell concentration is low. Guided by measurements of positive and negative strands in CVB3- infected cells (STAR Methods), we selected a threshold of 25 3D^pol^ molecules, which triggers at ∼2.5 hr after infection with 10 PFU and coincides with the first translational burst of an infectious virion (Boersma et al., 2020). This threshold marks the early onset of the deterministic, continuous regime (3D^pol^ counting noise = 20%) and precedes the 4–5-hr time point when VROs become ultrastructurally observable (Limpens et al., 2011).

To validate the parameterization of the replication module, we sought robust, sensitive, and absolute measurements of positive and negative strands during infection. We devised a tagged quantitative PCR (qPCR) assay, which avoids false priming within the CVB3 genome by using a biotinylated strand-specific primer and streptavidin pulldown before quantification (STAR Methods) (Bessaud et al., 2008; Singh et al., 2019). Using purified standards, the assay was sensitive to ∼1000 CVB3 copies per 250-cell reaction and linear over at least five decades (Figures S3B and S3C). After specifying a VRO transition consistent with these data, the model achieved excellent quantitative agreement with the absolute estimates of positive–negative strands (Figures 3C and S3D), and the module was deemed valid. Results were comparable if defective RNA genomes assembled as polysomes in the model or were further subject to nonsense-mediated decay (Figures S3E and S3F). Future iterations will consider the de novo generation of defective genomes during replication (Schulte et al., 2015).

Model–experiment concordance required the particle-to-PFU ratio described earlier and a quantification of the positive-to-negative strand ratio in released particles (1790; Figure S3B). Notably, we also assumed that replicative intermediates—partial RNA–RNA hybrids of positive and negative strand—were not detected by the tagged qPCR method. Upon cell lysis, collapsed replicative intermediates would be difficult to denature fully for biotinylated priming without thermally degrading the RNA itself. We tested the assumption by mixing 10^5^ copies of purified strand with increasing amounts of the complementary strand per 250-cell reaction and measuring both with the stranded assay (STAR Methods). Although copy numbers were accurate for many mixtures, some limiting ratios were irreproducible or grossly underestimated (Figure 3D). Such strand competition is probably even more severe in cells, where replicating strands are already nearby their complementary template (Novak and Kirkegaard, 1991).

There are experimental workarounds to accessing collapsed replicative intermediates, but their detection efficiency is difficult to determine (Hohenadl et al., 1991; Novak and Kirkegaard, 1991; Tam and Messner, 1999). Using the model at 10 PFU, we simulated the positive-to-negative strand ratio predicted when replicative intermediates were incorporated in the calculation at different efficiencies (STAR Methods). Replicative intermediates, even at 10% detection efficiency, profoundly altered the calculated ratio and its dynamic trajectory (Figure 3E). Literature-derived ratios of 50:1 to 100:1 were recapitulated in the model with detection efficiencies of 10–30% (Hohenadl et al., 1991; Leveque et al., 2012; Novak and Kirkegaard, 1991). At 100% detection efficiency, the model predicted lower ratios near 15:1. Lower positive-to-negative strand ratios have been observed in single-cell enterovirus assays involving large dilutions of a cell extract after lysis that may disfavor collapse of replicative intermediates (Schulte and Andino, 2014). Taken together, we conclude that the replication module is consistent with internal measurements as well as a range of observations in the literature.

### Encapsidation must coordinate the kinetics of enteroviral protein synthesis, recruitment, and self-assembly

Besides 3D^pol^, positive-strand translation also yields equal numbers of enteroviral proteinases, hydrophobic membrane-interacting proteins, and structural proteins that form the viral capsid (Figure 1A). We simplified by defining a single, lumped protease with the substrate specificity of both 2A^pro^ and 3C^pro^ (STAR Methods). Likewise, membrane-associated 3D^pol^ in the model considers 3D^pol^ together with the tightly interacting hydrophobic protein, 3AB (Figure 3A) (Xiang et al., 1998). We grouped the remaining hydrophobic proteins into a single exemplar, 2C^ATPase^, because of its direct interaction with the VP3 structural protein (Liu et al., 2010). VP3 assembles instantaneously with VP0 and VP1 to form a VP0–VP1–VP3 protomer (Jiang et al., 2014). We balanced molecular detail and complexity by modeling structural proteins as the 14S intermediate pentamer [(VP0–VP1–VP3)_5_], one fifth of which was generated with each polyprotein matured (Figure 4A). Together, these simplifications decompose the mature CVB3 polypeptide into four species—protease, 3D^pol^, 2C^ATPase^, and pentamer—that should balance stoichiometrically under limiting conditions (Figure S4A and S4B).

**Figure 4.**
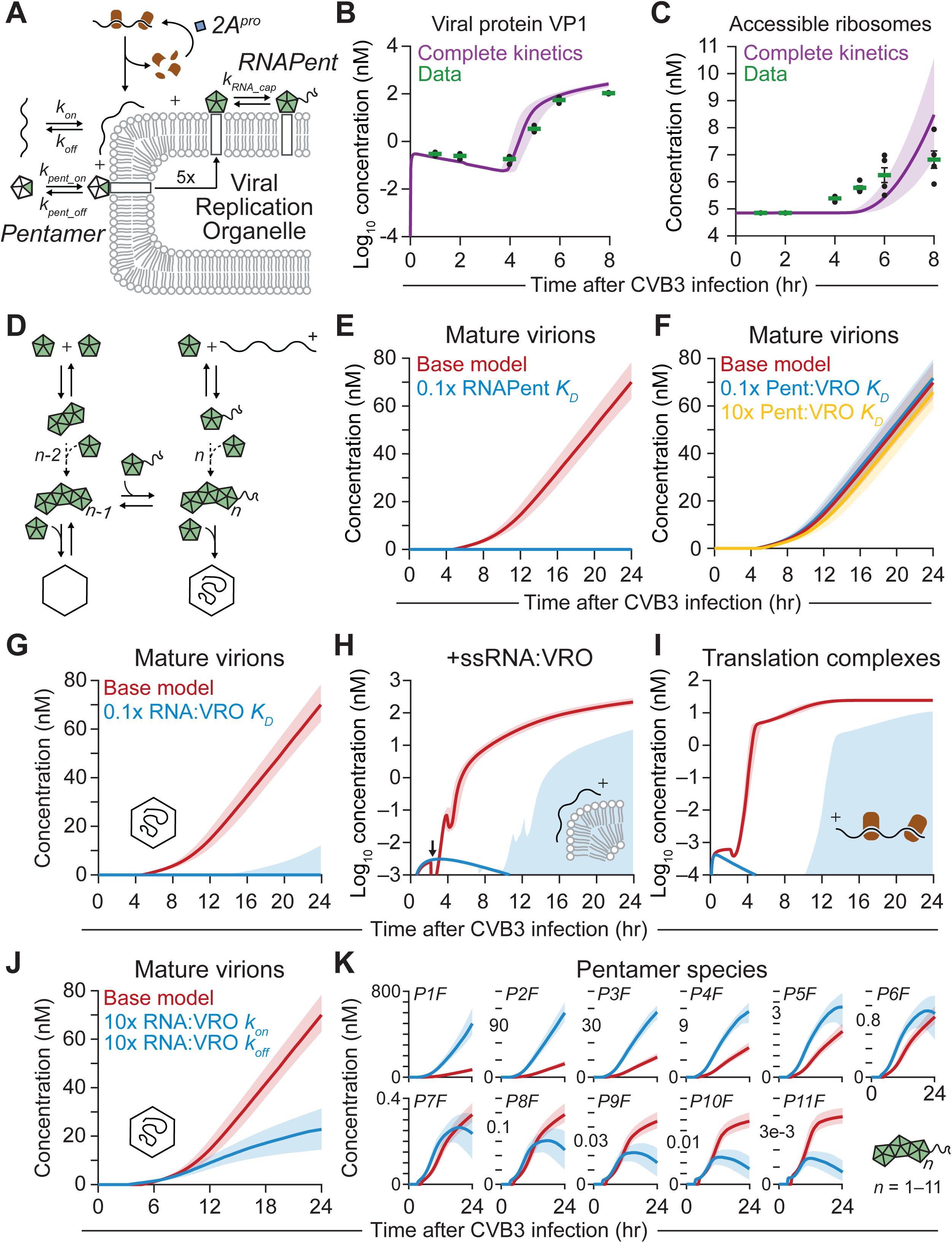
A Goldilocks zone for enteroviral encapsidation. (A) Overview schematic of the CVB3 encapsidation module on the viral replication organelle (VRO). After viral protein translation (aided by *2A^pro^* cleavage of eIF4G), viral capsid pentamers are recruited to VROs by binding 2C^ATPase^ (white box). At the VRO, pentamers associate with one another and with positive-strand genomes to form RNA-pentamer assemblies (*RNAPent*). Rate parameters investigated by perturbation are shown. (B and C) Complete kinetics captures the dynamics of viral protein VP1 expression and accessible ribosomes (estimated from viral protease cleavage of host eIF4G). Predictions (purple) were compared to data (green) obtained by quantitative immunoblotting. Data are shown as the geometric mean ± log-transformed standard error of *n* = 4 biological replicates of AC16-CAR cells infected at MOI = 10 for the indicated times. Population-level simulations are showing in Figure S4C and S4D. (D) Overview schematic of the CVB3 encapsidation module showing viral capsid assembly. Capsids assemble one pentamer at a time with or without a positive-strand genome. Filled virions (lower right) are formed irreversibly. (E) Increasing RNA-Pentamer affinity prevents virion maturation. Virion production in the base model (red) compared to when the RNA-pentamer affinity is increased (blue) by reducing its contact dissociation constant (RNAPent *K_D_*) from 1 mM to 100 µM (STAR Methods). (F) Model simulations are not sensitive to changes in pentamer recruitment to the VRO via 2C^ATPase^ binding. Virion production in the base model (red) compared to when pentamer-2C^ATPase^ affinity is either increased (blue) or decreased (yellow) by reducing or increasing its apparent dissociation constant (Pent:VRO *K_D_*) from 100 nM to 10 nM (blue) or 1 µM (yellow). (G–I) RNA interactions with the VRO must be balanced. (G) Virion production, (H) positive-strand RNA genomes at the VRO (+ssRNA:VRO), and (I) translation complexes in the base model (red) compared to when RNA exchange rates with the VRO were biased (blue) by decreasing *k_off_* to from 1 hr^-1^ to 0.1 hr^-1^. Arrow in (H) indicates premature recruitment of positive-strand RNA genomes to the VRO when *k_off_* = 0.1 hr^-1^. Some lognormally sampled parameter sets give rise to the late formation of +ssRNA:VRO, productive translation complexes, and mature virions (blue shading). (J and K) Kinetics of RNA-VRO interactions are important for sustained virion production. (J) Virion production and (K) pentamer states in the base model (red) compared to when RNA association and dissociation rates with the VRO are both increased tenfold (blue). PnF, intermediate filled capsid state of *n* pentamers and one positive-strand genome. For (B), (C), and (E–K), predictions are shown as the median simulation ± 90% nonparametric confidence interval from *n* = 100 simulations of single-cell infections at 10 PFU with a parameter coefficient of variation of 5%. See also Figure S4.

One of the earliest consequences of enteroviral proteinase maturation is the cleavage of the host-cell eukaryotic initiation factor, eIF4G, by 2A^pro^ (Etchison et al., 1982). eIF4G cleavage prevents ribosomes from initiating cap-dependent translation, thereby favoring the cap-independent translation of the virus (Figure 4A) (Darnell and Levintow, 1960; Fernandez-Munoz and Darnell, 1976). We modeled the shutoff of host-cell translation and theft of ribosomes by incorporating a protease-catalyzed conversion of ribosomes from inaccessible (cap-dependent) to accessible for CVB3 (STAR Methods). The full model showed good agreement with the relative dynamics of eIF4G cleavage and the synthesis of VP1 capsid protein (Figures 4B, 4C, S4C, and S4D), suggesting accurate encoding of the precursors to encapsidation.

Viral pentamers were not placed immediately on VROs like 3D^pol^ but instead were recruited to membranes by their interaction with available 2C^ATPase^ (Figure 4A; STAR Methods) (Liu et al., 2010). As neither kinetic nor equilibrium constants are available for the VP3–2C^ATPase^ interaction, we began with rate parameters that were realistic and consistent with a nominal affinity of 100 nM (Table S1). For viral RNA of either strand, there are multiple possible binding partners (Lyle et al., 2002), and thus we modeled VRO recruitment as a passive exchange between cytoplasmic and VRO compartments (STAR Methods). Considering the VRO concentrating effect and assuming a representative RNA-protein association rate of 25 nM^-1^hr^-1^ (Gleitsman et al., 2017), an exchange rate of 1 hr^-1^ equates to an effective membrane affinity of ∼125 nM. Sensitivity of the model to these approximations would be analyzed after defining the encapsidation steps that yield mature virions.

Encapsidation of enteroviral RNA arises from a series of individually weak interactions (∼1 mM contact affinity) that multiplicatively contribute to the final “closed” virion (Li et al., 2012), which we assumed to be irreversible (Figure 4D and Table S1; STAR Methods). Capsid self-assembly can also take place without viral RNA. We simplified the combinatorics by assuming that assembly occurs one pentamer at a time through reversible additions of pentamers that are either unbound or bound to RNA (STAR Methods). In this formalism, intermediate pentamer states can arise from multiple paths. For example, an empty state of five pentamers (*P5Empty*) can result from the addition of a pentamer to *P4Empty*, the loss of a pentamer from *P6Empty*, or the loss of RNA-bound pentamer from *P6Filled*. The disassembly of filled intermediates was assumed to release RNA with decreasing probability as the number of pentamers in the intermediate increased. *P6Filled*, for instance, disassembles to *P5Filled* + free pentamer ⅚of the time and *P5Empty* + RNA-containing pentamer ⅙of the time. The detailed encoding of encapsidation enables the module to keep track of the discrete steps to virion production and identify stoichiometric depletions of precursors or pentamer states if they occur.

To illustrate the importance of weak and balanced interactions for encapsidation, we increased the contact affinity between RNA and pentamer from 1 mM to 100 µM and found that the slight imbalance completely blocked encapsidation (Figure 4E). Pentamers recruited to the VRO were fully sequestered by the increasing abundance of positive strands, depleting the free pentamer pool available for higher-order assembly of capsids. Similar kinetic-trapping mechanisms dominated when pentamer contact affinities were tighter than 10 µM. By comparison, the module was not as sensitive to the VP3–2C^ATPase^ interaction strength that recruits pentamer to VROs, with virtually no change observed when the affinity was altered tenfold (Figure 4F and Table S1). Unrealistic delays in encapsidation did not occur until the affinity was reduced from 100 nM to 1–10 mM, suggesting that alanine mutants of 2C^ATPase^ disrupting encapsidation must severely hinder interactions with pentamer (Wang et al., 2012).

In contrast to pentamer recruitment, we found that kinetic interactions between viral RNA and the VRO surface were critical. Increasing the effective membrane affinity (tenfold decrease in *k_off_* and thus *K_D_*; Figure 4A) largely blocked capsid formation by driving RNA prematurely to the VRO at the expense of translation complexes needed to generate sufficient 3D^pol^ for replication (Figures 4G–I). Even with the same equilibrium affinity, faster exchange was problematic for effective encapsidation. We noted virion production tailed away when on–off rates were both increased tenfold (Figure 4J). The shorter residence time at the VRO led to large increases in early pentamer species (*P1Filled* to *P6Filled*) at the expense of later filled pentamers needed for sustained virion production (*P9Filled* to *P11Filled*; Figure 4K). The model thus provides a quantitative rationale for the multiple RNA-binding interactions at the membrane surface (Jiang et al., 2014). Overall, the encapsidation module finishes the viral life cycle in a stoichiometrically consistent way and makes specific predictions about the critical kinetic and thermodynamic steps of self-assembly.

### Loss of type I interferon signaling coincides with degradation of MAVS during CVB3 infection

With the viral life-cycle modules encoded, we returned to host-cell signaling responses in search of pathways engaged during an acute CVB3 infection. The CVB3 proteinase 3C^pro^ can block inflammatory signaling by cleaving the NF-κB inhibitor, IκBα, and converting it to a stably-associated, non-degradable form (Zaragoza et al., 2006). In AC16-CAR cells, however, we detected canonical degradation of IκBα and induction of NF-κB target genes a few hours after CVB3 infection (Figures 5A and 5B). We next considered the type I interferon signaling pathway, which is robustly triggered by cytosolic dsCVB3 in these cells (Figure 1B). Although phosphorylation of the effector kinase TBK1 was detected transiently after infection, the antiviral signal was not reliably propagated to IRF3 or interferon signaling through phosphorylated STATs (Figures 5C and 5D). There was no induction of the *IFNA* locus, nor did we observe upregulation of any ISG profiled in cells infected with live virus (Figures 5E and S5A–E). The results suggested that CVB3 actively severs signal transmission from its dsRNA replicative intermediate to a productive type I interferon response.

**Figure 5.**
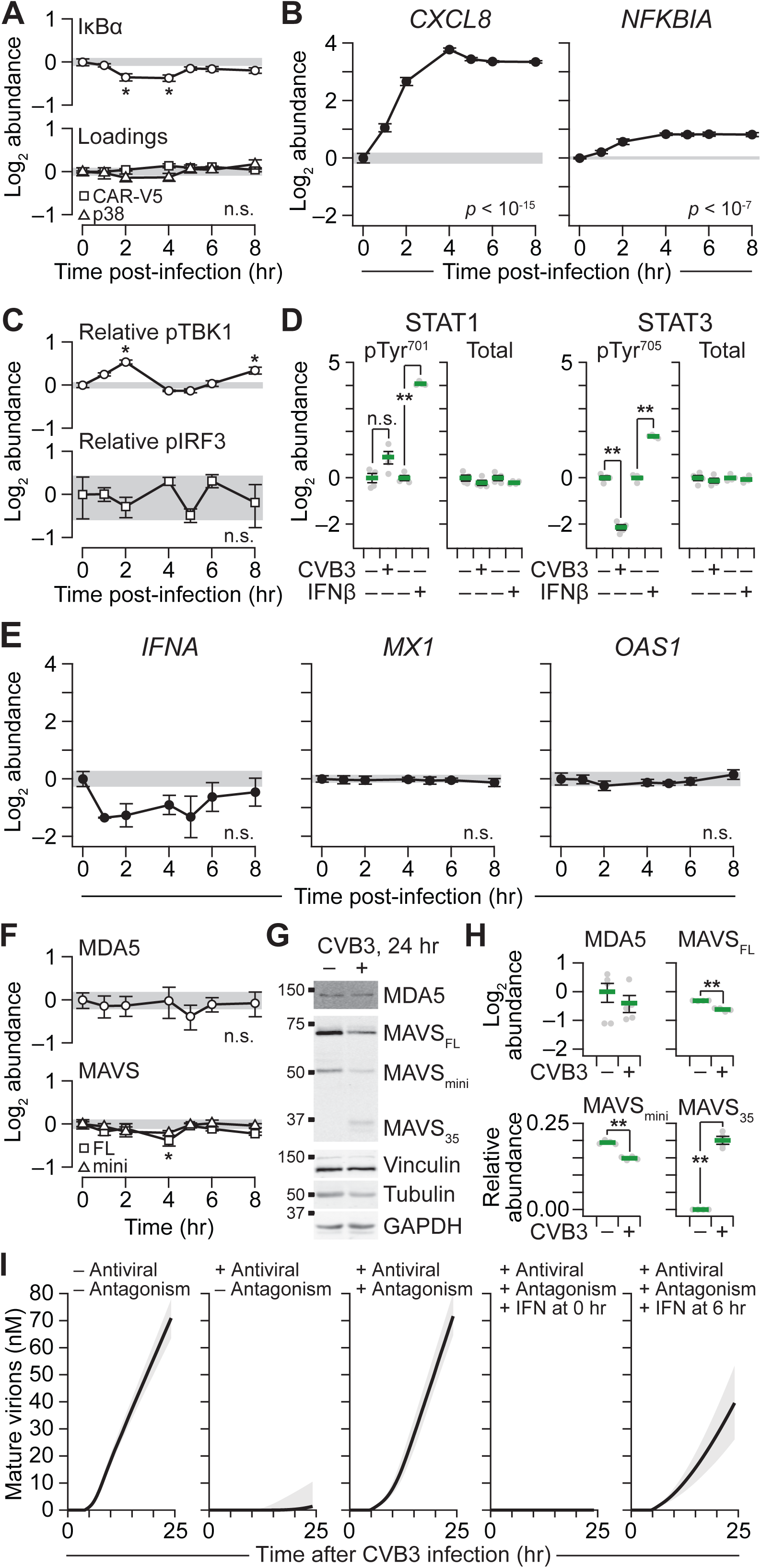
CVB3 partially dismantles antiviral signaling and the dsRNA transducer MAVS. (A and B) Acute CVB3 infection activates NF-κB signaling. (A) AC16-CAR cells were infected with CVB3 at MOI = 10 for the indicated times and immunoblotted for IκBα (A, upper) with ectopic CAR-V5, p38 (A, lower), vinculin, and tubulin used as loading controls. (B) Total RNA was collected at the indicated times and measured for the indicated NF-κB target genes with *HINT1*, *PRDX6*, and *GUSB* used as loading controls. (C–E) Early dsRNA sensing is not propagated to a detectable type I interferon response during acute CVB3 infection. (C) AC16-CAR cells were infected with CVB3 at MOI = 10 for the indicated times and immunoblotted for phosphorylated TBK1 (Ser^172^) relative to total TBK1 (Relative pTBK1) (C, upper) or phosphorylated IRF3 (Ser^396^) relative to total IRF3 (Relative pIRF3) (C, lower) with ectopic CAR-V5, p38, vinculin, and tubulin used as loading controls. (D) AC16-CAR cells were infected with CVB3 at MOI = 10 for 24 hours or treated with 30 ng/ml IFNβ for 30 minutes as a positive control and immunoblotted for phosphorylated STAT1 (pTyr^701^) and total STAT1 (D, left) or phosphorylated STAT3 (pTyr^705^) and total STAT3 (D, right) with vinculin, tubulin, and GAPDH (± CVB3) or actin, tubulin, and p38 (± IFNβ) used as loading controls. (E) Total RNA was collected at the indicated times and measured for the indicated ISGs with *HINT1*, *PRDX6*, and *GUSB* used as loading controls. (F–H) Acute CVB3 infection disrupts MAVS but not MDA5. (F) AC16-CAR cells were infected with CVB3 at MOI = 10 for the indicated times and immunoblotted for MDA5 (F, upper) or the full-length (FL) and mini-MAVS [mini; translated by leaky ribosomal scanning at Met^142^ (Brubaker et al., 2014)] with ectopic CAR-V5, p38, vinculin, and tubulin used as loading controls. (G) AC16-CAR cells were infected with CVB3 at MOI = 10 for 24 hours and immunoblotted for MDA5 or MAVS with vinculin, tubulin, and GAPDH used as loading controls. A 35 kDa MAVS cleavage product (MAVS_35_) is visible 24 hours after infection. (H) Immunoblot densitometry of replicated experiments described in (G). (I) Mature virion formation predicted by complete kinetics with or without innate antiviral sensing (antiviral), viral antagonism of antiviral sensing (antagonism), or supplemental interferon (IFN) added at the indicated times. Predictions are shown as the median simulation ± 90% nonparametric confidence interval from *n* = 100 simulations of single-cell infections at 10 PFU with a parameter coefficient of variation of 5%. Data are shown as the mean ± s.e.m. (A, C, D, F, H) or geometric mean ± log-transformed standard error (B, E) of *n* = 4 biological replicates. For (A–C), (E), and (F), time courses with significant alterations were assessed by one-way ANOVA (or, for MAVS_FL_ and MAVS_mini_, two-way ANOVA) with replication, and a single asterisk indicates *p* < 0.05 for individual time points compared to *t* = 0 hour (gray band) after Tukey post-hoc correction. For (D) and (H), a double asterisk indicates *p* < 10^-4^ by Student’s unpaired *t* test. See also Figure S5.

Both 3C^pro^ and 2A^pro^ are reported to target components of the innate dsRNA sensing and signal-transduction machinery (Feng et al., 2014; Mukherjee et al., 2011). Normally, long dsRNA is detected by MDA5, which associates with the mitochondrial transducer MAVS to drive surface polymerization that signals through the adaptor TRAF, the kinase TBK1, and the transcription factor IRF3 (Tan et al., 2018). Although MDA5 was not significantly altered in AC16-CAR cells during CVB3 infection, we noted a mid-infection dip in proteoforms of the mitochondrial transducer, MAVS (Figure 5F). Later post-infection times revealed a C-terminal fragment at 35 kDa (MAVS_35_), which comprised up to 20% of the endogenous MAVS protein at 24 hours; MDA5 was not clearly affected over the same timeframe (Figures 5G and 5H). The mid-infection decrease in full-length MAVS (MAVS_FL_) was concurrent with the loss of TBK1 phosphorylation and the explosive increase in CVB3 protein (Figures 4B, 5C, and 5F). A MAVS_35_ fragment implied that MAVS_FL_ was split in half, separating its N-terminal oligomerization domain from the C-terminal mitochondrial–microsomal transmembrane domain (Esser-Nobis et al., 2020; Seth et al., 2005). Even slight abundance shifts in MAVS proteoforms can alter the propensity of the pathway to activate (Brubaker et al., 2014; Qi et al., 2017). Together, the data suggested that a CVB3-derived protein antagonizes the dsRNA-mediated interferon response by cleaving an essential transducer (Sun et al., 2006).

We elaborated the infection model with a lumped interferon response, which pairs a CVB3 sensing-and-transduction mechanism with a single, pleiotropic ISG effector (*ISGs*, Figure 1A; STAR Methods). The joint sensor–transducer recognizes dsRNA and sigmoidally induces the ISG effector, streamlining the native regulation by the MDA5–MAVS–TRAF–TBK1–IRF3 pathway (Wu and Chen, 2014). The ISG effector impedes CVB3 infection at three points in the life cycle. First, the effector accelerates CVB3 RNA turnover, modeling the activity of oligoadenylate synthetases and RNAse L (Figures 1A and 1B) (Schwartz and Conn, 2019). Second, the effector blocks CVB3 polyprotein synthesis to capture dsRNA recognition and translational inhibition by the PKR pathway (STAR Methods) (Pindel and Sadler, 2011). Third, the effector incorporates interferon-stimulated mechanisms for disabling CVB3 proteinases, such as ISGylation of 2A^pro^ and PARP9–DTX3L-mediated ubiquitylation of 3C^pro^ (Rahnefeld et al., 2014; Zhang et al., 2015). The three antiviral effects were encoded as hyperbolic feedbacks on the CVB3 RNA degradation rates, the formation rate of CVB3 translation complexes, and the effective production rate of active 2A^pro^–3C^pro^ respectively (Figure 1A; STAR Methods). The active 2A^pro^–3C^pro^ pool reciprocally feeds back on the sensor–transducer in the model to hyperbolically limit the maximum induction rate of the ISG effector in response to dsRNA (Figure 1A; STAR Methods). The interlinking of these four negative feedbacks provides a compact abstraction of the multifaceted antagonism between enteroviruses and type I interferon signaling.

The feedback parameters of the lumped interferon response are phenomenological and not directly measurable. Nevertheless, the feedback architecture places strong qualitative constraints on their relative potencies:

1. AC16-CAR cells are capable of mounting an interferon response to CVB3 dsRNA (Figure 1B). However, the response is blocked during an active infection, presumably through protease-mediated antagonism of dsRNA sensing and cleavage of MAVS (Figure 5E–H). The feedback potency of the 2A^pro^–3C^pro^ pool must therefore be potent enough to degrade dsRNA sensing in the model during an active infection.
2. Early stimulation of AC16-CAR cells with interferons significantly impedes CVB3 propagation (Shah et al., 2017), suggesting that an endogenous interferon response would be sufficient if it were fully mobilized and not disrupted by viral proteases. Thus, the modeled type I interferon response should thwart the virus in simulations where dsRNA sensing–transduction is fully intact.
3. At later times during CVB3 infection, the addition of interferons becomes ineffective (Figure S5F). The 6–8 hr time window indicates that antiviral mechanisms become ineffective after the virus has passed a critical “point of no return” in its life cycle.

Using these constraints, we identified feedback weights with half-maximal effective concentrations (EC50s) that conferred the specified qualitative behavior (Figure 5I; STAR Methods). The outcome of infection was robust when these heuristic EC50 values were varied individually over a severalfold range (Figure S5G). Most EC50 values were in the nM range (5– 20 nM) except for protease antagonism of dsRNA sensing–transduction, which needed to be much lower (1 pM). Such potency could be achieved if the relevant protease is locally complexed with the dsRNA sensor–transducer that is cleaved. For MAVS cleavage, a plausible candidate is the precursor to 3C^pro^ and 3D^pol^, 3CD^pro^, which has both protease activity and nucleotide-binding activity for replication (Franco et al., 2005). The importance of potently antagonizing ISG induction is corroborated by a recent computational model of Dengue virus (Zitzmann et al., 2020). Combining the enteroviral life cycle with host-cell feedbacks created a rich dynamical system to mine for predicted behavior that was non-intuitive and possibly important for pathogenesis.

### A cleavage-resistant MAVS mutant shows enhanced antiviral activity upon delayed stimulation with paracrine interferon

We used the model to examine viral outcomes to delayed addition of interferon (Figure S5F). The scenario mimics a setting of paracrine antiviral signaling, where an infected cell is warned of a local infection by an interferon-secreting cell nearby. The earlier this paracrine warning signal is received, the more effective the stimulated interferon response can be at inhibiting viral propagation. We reasoned that the timed addition of interferon would uncover nonobvious combinatorial sensitivities in the network that could be tested experimentally. Using mature virions as the readout, we screened a five-tiered range of potencies for the three interferon feedbacks and viral-protease antagonism (5^4^ = 625 simulations), and each simulation was run with or without interferon at five different times after infection [5^4^(1+5) = 3750 simulations]. Many of the perturbations affected mature virions monotonically with effects that superposed when combined. However, the computational screen revealed a sensitivity to viral-protease antagonism that was dependent on delayed addition of interferons at intermediate times post-infection (Figures 6A and 6B). If interferon was supplemented early (t ∼ 4 hr), viral propagation was dampened to a basal level regardless of the extent of antagonism. However, when viral-protease antagonism was reduced fivefold, the dampening persisted even when interferon was supplemented later (t ∼ 6 hr). At this time, the base model cannot maintain peak ISG production because of excess viral protease but the resistant variant is still maximally fighting the infection (Figures S6A and S6B). The testable prediction was that perturbing sensor–transducer degradation by viral proteases would have analogous time-dependent effects when infected cells received supplemental interferons.

**Figure 6.**
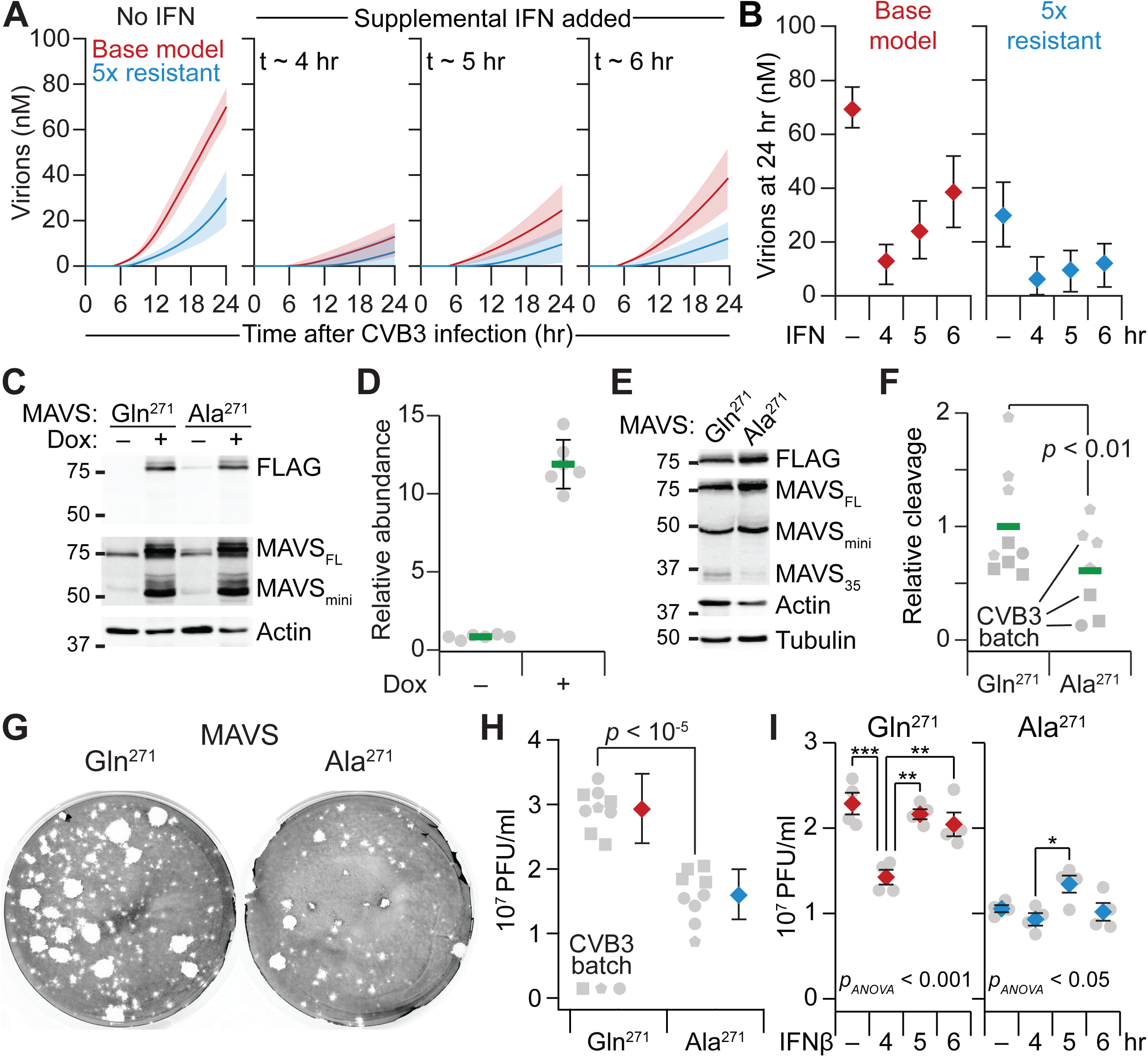
MAVS is a sensitive locus for CVB3 susceptibility in the host-cell network. (A and B) CVB3 virion production is modulated nonlinearly by host-cell resistance and time-delayed supplementation of interferon. Virion output from complete kinetics with (blue) or without (red) fivefold altered resistance to viral proteinases. Negative feedback from the viral proteinases on the viral dsRNA sensor–transducer was decreased to mimic an increase in host-cell resistance. Supplemental interferon (IFN) was simulated at the indicated times. The 24-hour terminal endpoint of the simulations in (A) are summarized in (B). Predictions are shown as the median simulation ± 90% nonparametric confidence interval from *n* = 100 simulations of single-cell infections at 10 PFU with a parameter coefficient of variation of 5%. (C and D) Inducible ectopic expression of MAVS alleles. (C) AC16-CAR cells stably transduced with doxycycline (Dox)-regulated 3xFLAG-tagged Gln^271^ MAVS or Ala^271^ MAVS were treated with 1 µg/ml Dox for 24 hours and immunoblotted for FLAG and MAVS with actin used as a loading control. The image gamma was adjusted for MAVS (gamma = 4) to show the endogenous full-length MAVS (MAVS_FL_) and mini-MAVS (MAVS_mini_) at the same exposure as the induced constructs. (D) Immunoblot densitometry of induced ectopic MAVS relative to endogenous MAVS_FL_ + MAVS_mini_ in AC16-CAR cells treated with or without Dox. Data are shown as mean ± s.d. of *n* = 6 different MAVS alleles used in this work (see also Figure 7; STAR Methods). (E and F) CVB3-induced MAVS cleavage is reduced in cells expressing the Ala^271^ mutant. (E) AC16-CAR cells stably transduced with inducible Gln^271^ MAVS or Ala^271^ MAVS were induced with Dox for 24 hours, then infected with CVB3 at MOI = 5 for 24 hours and immunoblotted for FLAG or MAVS with actin and tubulin used as loading controls. MAVS_FL_, MAVS_mini_, and the 35 kDa MAVS cleavage product (MAVS_35_) are indicated. (F) Immunoblot densitometry of replicated experiments described in (E) using *n* = 3 different CVB3 batches and 1–4 biological replicates per batch. (G and H) CVB3-induced virion release is reduced in cells expressing Ala^271^ MAVS. (G) Representative plaque assay for infectious virion release in the conditioned medium from AC16-CAR cells after induction of Gln^271^ MAVS or Ala^271^ MAVS and infection with CVB3 at MOI = 5 for 24 hours. (H) Quantification of plaque-forming units (PFU) from *n* = 3 different CVB3 batches and 1–4 biological replicates per batch. Data are summarized as PFU/ml ± 95% Poisson confidence intervals based on the mean or the observation. (I) Ala^271^ MAVS sustains its antiviral potency upon delayed addition of beta-interferon (IFNβ). After induction of Gln^271^ MAVS or Ala^271^ MAVS, AC16-CAR cells were infected with CVB3 at MOI = 10 for 24 hours, with 30 ng/ml IFNβ added at the indicated times after the start of CVB3 infection. Data are shown as the mean ± s.e.m. of *n* = 4 biological replicates, and differences across conditions was assessed by one-way ANOVA (*p_ANOVA_*) with Tukey’s posthoc test for individual differences: * *p* < 0.05, ** *p* < 0.01, *** *p* < 0.001. The indicated reductions in viral titers between the two genotypes are qualitatively similar to a second Ala^271^-harboring MAVS allele (Ala^148^Ala^271^) shown in Figure S6D. For (F) and (H), the difference between MAVS genotypes was assessed by replicated two-way ANOVA with MAVS genotype and CVB3 batch as factors. See also Figure S6.

To test this prediction, we returned to the CVB3-induced cleavage of MAVS observed earlier (Figures 5G and 5H). MAVS can be cleaved by 3C^pro^ at a Gln-Ala cleavage site between positions 148 and 149 (Mukherjee et al., 2011). However, such a cleavage product would be electrophoretically indistinguishable from MAVS_mini_, an oligomerization-deficient proteoform that is translated by leaky ribosomal scanning at Met^142^ (Brubaker et al., 2014). The site’s importance for CVB3 infectivity is also unclear, because Ala^149^ in MAVS is undergoing rapid positive selection in primates (Patel et al., 2012). Chimpanzees (Val^149^) and African green monkeys (Arg^149^) both have substitutions that should prevent cleavage by 3C^pro^ (Blom et al., 1996), but coxsackievirus infections have been documented in both species (Nielsen et al., 2012; Takada et al., 1968). We considered alternative sites that would be more consistent with the observed MAVS_35_ product and identified a Gln-Gly cleavage site between positions 271 and 272. The site is included in a six amino-acid insertion distinguishing old-world monkeys/hominoids from new-world monkeys, which are seronegative for coxsackievirus (Deinhardt et al., 1967). The phylogenetics and observed cleavage pattern of MAVS built a stronger case for Gln^271^ as an important site for sensor–transducer degradation during CVB3 infection.

Using a human MAVS template derived from a widely cited IMAGE clone (Meylan et al., 2005; Seth et al., 2005), we cloned a doxycycline-inducible, FLAG-tagged MAVS with or without Gln^271^ substituted for alanine (Ala^271^; STAR Methods). Lentiviral transduction and selection of AC16-CAR cells yielded lines with comparably inducible ectopic MAVS alleles, which could be compared to one another (Figures 6C and 6D). When cells were induced and then infected with CVB3, we found that generation of MAVS_35_ was significantly reduced in the Ala^271^ line compared to the Gln^271^ line for multiple viral stocks (Figures 6E and 6F). We attributed the residual MAVS_35_ in the Ala^271^ line to cleavage of endogenous MAVS at Gln^271^. Thus, CVB3 antagonizes antiviral signal transduction by cleaving MAVS at Gln^271^, and the Ala^271^ line approximates a state of partial resistance as simulated by the complete kinetic model.

To evaluate the impact of MAVS cleavage on viral propagation, we collected medium from cells treated with CVB3 for 24 hours and titered infectious virions by plaque assay (STAR Methods). Titers from the Ala^271^ line were significantly lower relative to the Gln^271^ line for multiple CVB3 stocks (Figures 6G and 6H). Biological error was usually close to the intrinsic counting noise of a plaque assay (Figures S6C and S6D), with the spread of biological replicates comparable to Poisson intervals about the mean (Figure 6H). We also kept in mind the increased uncertainty of the interferon-supplemented base model (Figure 6B, left) when comparing predictions to experiments. The Gln^271^ and Ala^271^ lines were next used to test for differences in the time-dependent effect of paracrine interferon signaling predicted by the model. We found that interferon-β treatment at 4 hours after infection significantly reduced the infectivity in the Gln^271^ line, as paracrine stimulation offset the MAVS genotype (Figure 6I, left). At 5–6 hours, however, interferon-β lost efficacy in the Gln^271^ line, returning to viral titers that were observed without paracrine stimulation. The Ala^271^ line, by contrast, showed little, if any, time-dependent effect for interferon-β supplementation (Figure 6I, right). The Ala^271^ mutant agreed even more closely with model predictions when using an inducible, cleavage-resistant MAVS allele containing a compound mutation at the other site reportedly cleaved by 3C^pro^ (Ala^148^Ala^271^; Figure S6E) (Mukherjee et al., 2011). We conclude from these experiments that the overall importance of CVB3-proteinase susceptibility for MAVS is strongly modulated by paracrine interferon, consistent with the predictions of complete kinetics.

### Enteroviral proteinase cleavage of MAVS is redirected by a prevalent human polymorphism

The IMAGE clone used to construct the MAVS alleles is valid but contains three common polymorphisms that deviate from the reference protein sequence of humans and other mammals: Q93E (rs17857295; 25% aggregated allele frequency), Q198K (rs7262903; 16%), and S409F (rs7269320; 16%) (Phan et al., 2020). The Q198K polymorphism is considered functionally neutral (Pothlichet et al., 2011), and none of these variants alter the mitochondrial localization of MAVS (Xing et al., 2016). However, the substitution at position 93 was relevant, because Gln^93^ in the reference sequence creates the possibility of an additional site for cleavage by 3C^pro^ (Figure 7A) (Blom et al., 1996). Like Gln^271^, cleavage at Gln^93^ would separate the oligomerization domain (CARD) of MAVS and thereby prevent signal transmission from MDA5 to TRAFs on the mitochondrial surface (Figure 7A).

**Figure 7.**
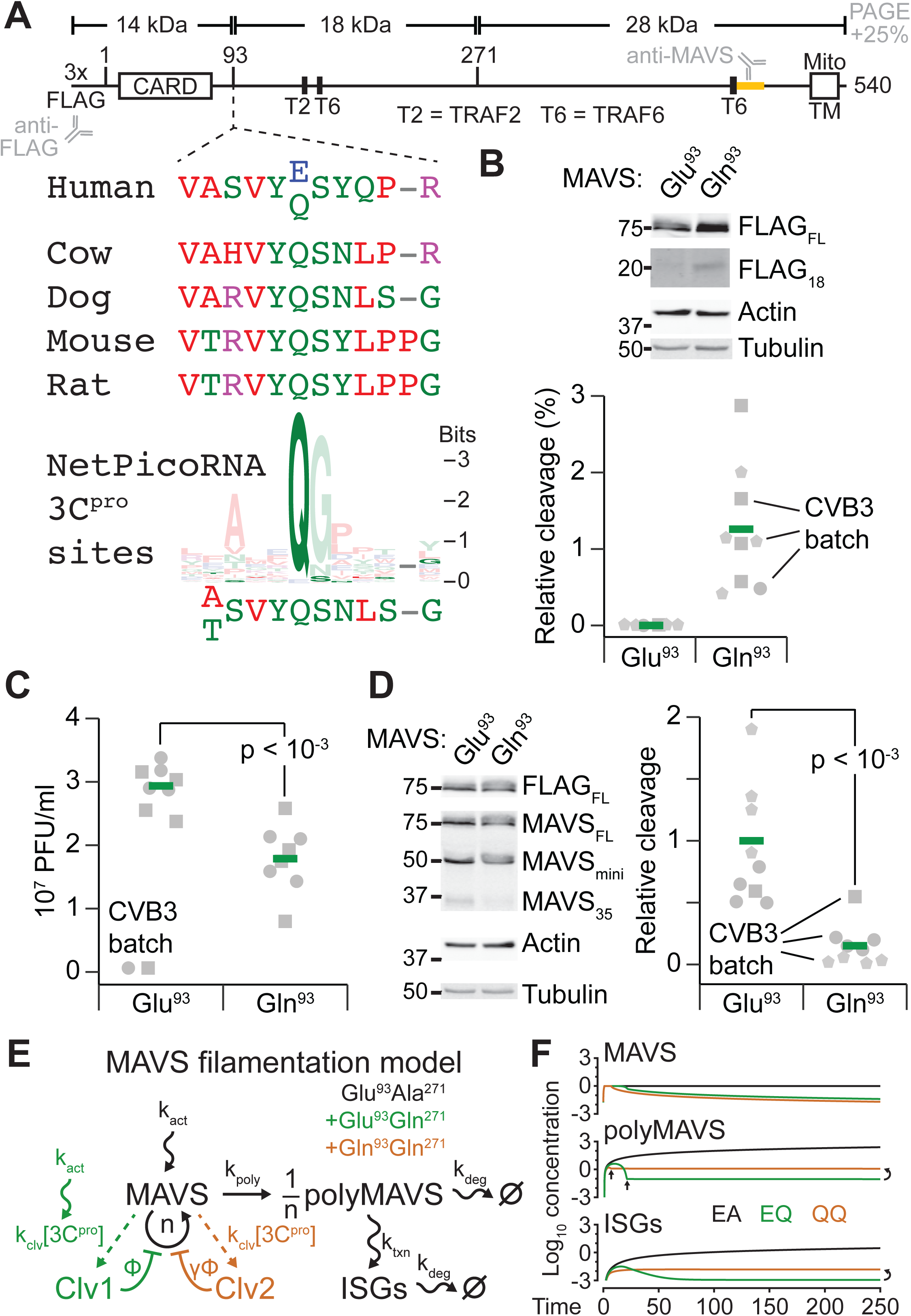
MAVS is a sensitive locus for CVB3 susceptibility in the human population. (A) Sequence and domain architecture of MAVS. Above: predicted molecular weights of MAVS cleaved at the indicated positions—MAVS is acidic and separates on an SDS-polyacrylamide gel (PAGE) with an electrophoretic mobility ∼25% larger than the predicted molecular weights listed. The oligomerization domain (CARD) and mitochondrial transmembrane (Mito TM) domains are indicated along with the recruitment sites for TRAF2 (T2) and TRAF6 (T6). The N-terminal epitope tag is indicated as well as the approximate peptide epitope of the anti-MAVS antibody (yellow). Below: flanking amino acids around position 93 of human MAVS, which has a Glu/Gln polymorphism. Gln^93^ is widely conserved in mammals. The sequence logo for enteroviral 3C^pro^ was re-derived from the enteroviral cleavage sites analyzed by NetPicoRNA (STAR Methods) (Blom et al., 1996). (B) Ectopic expression of Gln^93^ MAVS gives rise to an 18 kDa cleavage fragment. AC16-CAR cells stably transduced with inducible Glu^93^ MAVS or Gln^93^ MAVS were induced with doxycycline for 24 hours, then infected with CVB3 at MOI = 5 for 24 hours and immunoblotted for FLAG with actin and tubulin used as loading controls. The image gamma was changed for FLAG_18_ (gamma = 4). (bottom) Immunoblot densitometry of replicated experiments using *n* = 3 different CVB3 batches and 1–4 biological replicates per batch. The difference between MAVS genotypes was assessed by replicated two-way ANOVA with MAVS genotype and CVB3 batch as factors. (C) Ectopic Gln^93^ MAVS significantly reduces virion release compared to ectopic Glu^93^ MAVS. AC16-CAR cells stably transduced with inducible Glu^93^ MAVS or Gln^93^ MAVS were induced with Dox for 24 hours, then infected with CVB3 at MOI = 5 for 24 hours. Infectious virions in the conditioned medium were collected at the end of the 24-hour CVB3 infection. Quantification of plaque-forming units (PFU) is from *n* = 2 different CVB3 batches and four biological replicates per batch. The difference between MAVS genotypes was assessed by replicated two-way ANOVA with MAVS genotype and CVB3 batch as factors. (D) Ectopic expression of Gln^93^ MAVS reduces the 35 kDa cleavage fragment. AC16-CAR cells stably transduced with inducible Glu^93^ MAVS or Gln^93^ MAVS were induced with Dox for 24 hours, then infected with CVB3 at MOI = 5 for 24 hours and immunoblotted for FLAG and MAVS with actin and tubulin used as loading controls (left). Immunoblot densitometry of replicated experiments using *n* = 3 different CVB3 batches and 1–4 biological replicates per batch (right). The difference between MAVS genotypes was assessed by replicated two-way ANOVA with MAVS genotype and CVB3 batch as factors. (E) State-based model of MAVS self-assembly and cleavage by 3C^pro^. The core model for the Glu^93^Ala^271^ mutant (black) is elaborated with one or two 3C^pro^-catalyzed cleavage reactions for the Glu^93^Gln^271^ MAVS (green) or the Gln^93^Gln^271^ polymorphism (brown), respectively. (F) Simulated trajectories for the three MAVS alleles: Glu^93^Ala^271^ (EA, black), Glu^93^Gln^271^ (EQ, green), and Gln^93^Gln^271^ (QQ, brown). Single-parameter sensitivity analysis for the time-integrated ISG profile is shown in Figure S7A, and the effect of MAVS genotypes in Figures S7B–D. See also Figure S7 and Note S1.

Considering the effect of the synthetic Ala^271^ mutant (Figures 6C–I), we reasoned that position 93 might represent a naturally variable site affecting CVB3 infectivity in the human population. We replaced Glu^93^ in the IMAGE clone sequence with Gln^93^ and established AC16-CAR lines with equivalently inducible expression (Figure 6D). Upon CVB3 infection of the Gln^93^ line, we detected a specific N-terminal MAVS fragment at ∼18 kDa, which was consistent with cleavage at position 93 (Figures 7A and 7B). Given that the Gln^93^ allele of MAVS harbors an additional 3C^pro^ site at Gln^271^, we anticipated that this line would be even more permissive to CVB3 infection than the Glu^93^ line. Unexpectedly, the opposite was true—the Gln^93^ line yielded significantly fewer infectious virions than the Glu^93^ line, despite twice as many 3C^pro^-targetable sites (Figure 7C). Moreover, the Gln^93^ allele of MAVS was cleaved much less extensively at position 271 in CVB3-infected cells, as indicated by the formation of MAVS_35_ (Figure 7D). The results indicated a coupling between Gln^93^ and Gln^271^, where two cleavage options for 3C^pro^ rendered MAVS more resistant to viral antagonism than one.

There are now detailed models of MAVS activation triggered by dsRNA (Schweinoch et al., 2020). However, before stipulating hierarchical or otherwise-special properties for different MAVS cleavages, we asked whether a separate, highly simplified model of MAVS regulation could be explanatory (Figure 7E). MAVS signaling occurs on the surface of mitochondria by polymeric self-assembly into filaments, which are nucleated by MDA5 binding to the dsRNA intermediates of CVB3 (Feng et al., 2012; Hou et al., 2011). For the model, we assumed that MAVS activation occurs at a constant rate (*k_act_*) and on the same time scale as the activation of 3C^pro^ (STAR Methods). Structural data suggest that polymeric MAVS (polyMAVS) filaments are large (up to *n* = 800 molecules) (Wu et al., 2013; Wu et al., 2014). We modeled polyMAVS formation as an instantaneous assembly given that it occurs on a two-dimensional surface, and signaling from polyMAVS to ISGs was lumped as a first-order “transcription” process (*k_txn_*). In cells, polyMAVS is degraded slowly by mitophagy (Qi et al., 2017), and we assumed a similar decay constant for ISGs (*k_deg_*; Figure 1C). These equations governed the behavior of the Glu^93^Ala^271^ allele that was assumed to be uncleavable by 3C^pro^.

For cleavable alleles, we appended one (Glu^93^Gln^271^) or two (Gln^93^Gln^271^) first-order pathways in which 3C^pro^ slowly splits MAVS (*k_clv_*) and removes it from the pool of monomers for polymerization (Clv1 or Clv2; Figure 7E). Either cleavage will remove the oligomerization domain from MAVS and render it unable to polymerize (Figure 7A). Recognizing that the remaining C-terminal truncations on the mitochondrial surface could further inhibit the polymerization of full-length MAVS (Qi et al., 2017), we allowed for negative feedback on the polymerization. In addition, we permitted unequal feedback between the two cleavage products for the Gln^93^Gln^271^ allele (Φ and γΦ), even though the simulations were largely insensitive to negative feedback overall (Figure S7A). When negative feedbacks are equal (Figure 7F) or absent (Note S1), the only difference between the simulations involving cleavage is that the polymerization-competent Gln^93^Gln^271^ is removed twice as quickly as Glu^93^Gln^271^.

In the filamentation model, all MAVS genotypes rapidly reach a pseudo-steady state as polyMAVS and ISGs begin to accumulate (Figure 7F). This trajectory is sustained for the Glu^93^Ala^271^ allele, because nothing abates the constant rate of MAVS activation (EA; Figure 7F). For the cleavable alleles, however, 3C^pro^ eventually reaches a concentration that depletes the reservoir of monomeric MAVS, causing polyMAVS to decline. When MAVS depletion reaches the point that polyMAVS formation is negligible compared to its mitophagic disposal, the kinetics of polyMAVS shift to a first-order decay with a long time constant (Note S1). For the Gln^93^Gln^271^ allele (QQ; Figure 7F), polyMAVS both decelerates and troughs twice as quickly, hence decaying slowly from a higher concentration of polyMAVS than that of the Glu^93^Gln^271^ allele (EQ; vertical arrows in Figure 7F). In this set of model parameters, the distinction is enough to change ISG kinetics from sustained to transient, but differences in ISG induction are preserved for a range of parameter sets (Figure S7A). The model illustrates how non-intuitive antiviral behavior arises when the kinetic competition between MAVS and 3C^pro^ is funneled through a surface-polymerization step that is highly nonlinear. If small differences in proteinase susceptibility are amplified by paracrine interferons (Figure 6), the minor-but-prevalent Glu^93^ allele may thus contribute to the individual severity of human enteroviral infections.

## DISCUSSION

This work examines the feasibility of complete kinetics for acute viral infections by leveraging the unsegmented genome of picornaviruses and 70+ years of enteroviral research in cells (Enders et al., 1949). We mathematically encoded the molecular pathways from viral binding and entry through the formation of mature virions. Interference by host-cell signaling was not ignored but superimposed as a set of feedbacks, which themselves were subject to increasing antagonism during viral progression. The resulting draft is consistent with measured parameters and other observations in the literature, as well as our own experiments. Collectively, the formalized mechanisms, computer simulations, and experimental results point to MAVS as a critical determinant of the enteroviral response in human cells.

### Generalization to other enteroviruses and cellular contexts

Organizing the complete kinetics of CVB3 into modules is advantageous (Figure 1A), because it will streamline adaptation to other enteroviruses. Within the genus, the most substantive differences lie in binding and entry. Poliovirus, for instance, uses a single cell-surface receptor with two binding affinities and delivers genome to the cytoplasm before it is fully endocytosed (Brandenburg et al., 2007; McDermott et al., 2000; Racaniello, 2013). Exchange of a poliovirus-specific delivery module may be sufficient to explain the accelerated kinetics of infection relative to CVB3.

For any enterovirus species, results from complete kinetics suggest that the particle-to-PFU ratio is a critical determinant for predicting host-cell tropism and susceptibility. Viruses with large ratios of defective particles will depend on excess cell-surface receptors to infect, which may not be true in all humans. Receptors for coxsackievirus decline considerably with age (Kunin, 1962), presumably to different extents among individuals. For CAR, the interquartile range of transcript expression is 1.7–5.6 TPM (4000–10,000 copies per cell) in the left ventricle (Consortium, 2015), which straddles the inferred threshold for productive infection with the CVB3 stocks used here (∼7000 copies per cell; Figures 2E and S2F). The delivery module can accommodate such host-cell variation in a scalable and principled way.

### Different VROs for different kinetic regimes of viral infection

All positive-strand RNA viruses reconfigure host-cell membranes to promote replication (Miller and Krijnse-Locker, 2008). However, the membrane rearrangements differ among viruses, and it is unclear whether function of the resulting VROs is the same. Complete kinetic modeling of CVB3 indicates that the main advantage conferred by enteroviral VROs is to accelerate replication biochemistry on membrane surfaces. The hydrophobic 3AB protein binds 3D^pol^ and supports its intrinsic propensity to form two-dimensional lattices (Lyle et al., 2002). In turn, VROs emerge right at the onset of exponential enterovirus replication (Limpens et al., 2011).

A computational model of flavivirus reached a different conclusion about the role of VROs (Binder et al., 2013). Parameter sensitivity of VROs in this model emphasized the importance of compartmentalizing RNA-dependent RNA polymerase and shielding the positive-strand template from degradation. The apparent discrepancy can be reconciled when considering the different rates of infection and replication between the two RNA-virus genera. Flaviviral polymerase associates >60-fold more slowly and synthesizes at one-third the rate of enteroviruses (Jin et al., 2011) (Table S1), with infections requiring multiple days to establish. The much slower time scales should increase the importance of sequestering flavivirus RNA with polymerase and away from host-cell ribonucleases. Indeed, flaviviral VROs assemble as membranous webs or invaginations that shield replication components from the host (Miller and Krijnse-Locker, 2008), supporting kinetics as a pressure for viral evolution.

### Expanding and revising complete kinetic models

We emphasize that the complete kinetics of CVB3 reported here is a draft that should be subject to future refinements. Many reaction parameters were drawn from literature on poliovirus (Table S1), which is more similar to certain A-type coxsackievirus strains than to CVB3 (Brown et al., 2003). Also, the physical interactions between viral RNA and the VRO surface were not specified with the same detail as elsewhere in the kinetic model. If the longer-lived 3CD^pro^ precursor confers specific RNA-binding properties to the membrane (Herold and Andino, 2001), the stoichiometry of this intermediate may need to be considered explicitly. The threshold for VROs was the one instance where a model parameter was “tuned” to data. Although uncertainty may persist around the switch to VRO-like surface behavior, future alternatives will be easy to vet because VRO initiation is such a critical transition in the simulations.

The feedbacks overlaid on the kinetic model could certainly be elaborated more deeply, but there are also advantages to lumping at this scale. For example, MAVS is cleaved at a different site (Gln^428^) by the mitochondrially localized 3ABC precursor of hepatitis A picornavirus (Yang et al., 2007). The net result, however, is the same as cleavage at Gln^93^ or Gln^271^— separation of the MAVS oligomerization domain from the mitochondrial surface where it can polymerize rapidly. A different challenge relates to incorporating the human MAVS alleles with different cleavage susceptibilities. Although the simplified MAVS regulatory model is intriguing, it is premature to incorporate as part of complete kinetics. The modeled step is but one in a pathway, using parameters that were nominal approximations of fast and slow processes. Even within this step, the true length of polymerized MAVS filaments in cells is actively debated (Hwang et al., 2019). Frankly, cell signaling is not really geared for complete kinetics like fast-acting RNA viruses are. For cell signaling, more appropriate are the abstractions and flexible system boundaries of models that are incomplete but useful (Janes and Lauffenburger, 2013).

### Implications for enteroviral disease

The Q93E polymorphism in MAVS occurs with a mutant allele frequency of ∼50% among individuals of East Asian ancestry (Phan et al., 2020), indicating a large proportion of homozygous individuals in these populations. Geographically, many Asian countries face widespread cyclical outbreaks of enterovirus infection, and the understanding of risk factors is incomplete (Koh et al., 2016). The MAVS regulatory model suggests that a pure Glu^93^ context behaves very differently from a mixture of Glu^93^ and Gln^93^ (Figures S7B–D). Based on the *MAVS* RNA-seq reads, we gleaned that AC16 cells are heterozygous for the Q93E allele. Our results argue for more-formal studies of CVB3 susceptibility in isogenic lines that investigate different genotypes of MAVS expressed at endogenous levels.

Pharmacologically, 3C^pro^ has long been recognized as a therapeutic target for enteroviral infections, with inhibitors reaching as far as Phase II (Becker et al., 2016; Hayden et al., 2003; Kim et al., 2012). The rationale for such compounds is to block maturation of the enteroviral polypeptide, but it can be challenging to target an intramolecular cleavage with sufficient potency in cells. Intermolecular 3C^pro^ activity toward MAVS provides a means for reappraisal, especially considering its variable cleavability in humans. According to the encoded viral feedback, a 10–20-fold shift in the EC50 for sensor–transducer degradation is sufficient to block a low-titer CVB3 infection (MOI = 1) entirely.

Complete kinetics is a tangible organizing principle for viruses with a limited gene repertoire and an acute mode of infection. The concept may seem a distant goal for large viruses with multiple unknown gene products. It is nonetheless a goal that can specify where operational paradigms are unsatisfactory. Overall, viruses are much more modular than the host cells they infect. Systems bioengineers should exploit this property to define the integrated parts lists that could one day be mixed and matched to propose complete kinetic models for viruses that have recently recombined (Janes et al., 2017; Li et al., 2020).

## ACKNOWLEDGMENTS

We thank Peter Kasson and Cameron Griffiths for comments regarding the manuscript, Emily Farber at the Center for Public Health Genomics for performing the RNA-seq, Matthew Sutcliffe for performing the RNA-seq alignments, Lixin Wang for recognizing the Q93E substitution in the original MAVS IMAGE clone, and Karin Klingel for reagents. This work was supported by The David and Lucile Packard Foundation (Grant 2009-34710, K.A.J.), a gift from the Salamone Family Foundation (K.A.J.), and the UVA Cardiovascular Training Grant (Grant T32-HL007284, A.J.S.).

## AUTHOR CONTRIBUTIONS

Conceptualization, A.B.L, A.J.S., C.M.S., E.G.G., M.S., and K.A.J.; Methodology, A.J.S., C.M.S., E.G.G., M.S., and K.A.J.; Software, A.B.L., C.M.S., M.S., and K.A.J.; Validation, C.A.B.; Formal Analysis, A.J.S., M.S., and K.A.J.; Investigation, A.B.L., A.J.S., C.M.S., E.G.G., M.S., and K.A.J.; Resources, A.B.L., A.J.S., C.M.S., C.A.B., M.S., and K.A.J.; Data Curation, A.B.L., A.J.S., C.A.B., M.S., and K.A.J.; Writing – Original Draft, A.B.L., A.J.S., C.M.S., C.A.B., M.S., and K.A.J.; Writing – Review & Editing, A.B.L., A.J.S., C.A.B., M.S., and K.A.J.; Visualization, A.J.S., M.S., and K.A.J.; Supervision, C.M.S., M.S., and K.A.J.; Project Administration, M.S. and K.A.J.; Funding Acquisition, A.J.S. and K.A.J.

## STAR METHODS

### RESOURCE AVAILABILITY

#### Lead Contact

Further information and requests for resources and reagents should be directed to and will be fulfilled by the Lead Contact, Kevin A. Janes.

#### Materials Availability

Plasmids related to this work are deposited with Addgene (#158628–158646).

#### Data and Code Availability

- RNA-seq source data have been deposited at the NCBI Gene Expression Omnibus and are publicly available under the accession number: GSE155312.
- MATLAB original code for the complete kinetic model of CVB3 is publicly available at https://github.com/JanesLab/CompleteKinetics-CVB3. MATLAB original code for the MAVS filamentation model is publicly available at https://github.com/JanesLab/MAVSfilamentation.
- The scripts used to generate the modeling figure subpanels reported in this paper are available at https://github.com/JanesLab/LopacinskiAB-CellSyst2021. Scripts were not used to generate the experimental figure subpanels reported in this paper.
- Any additional information required to reproduce this work is available from the Lead Contact.

### EXPERIMENTAL MODEL AND SUBJECT DETAILS

#### Cell lines

AC16 (Davidson et al., 2005) and AC16-CAR cells (Shah et al., 2017) were cultured in DMEM/F12 (Gibco) + 12.5% fetal bovine serum (Hyclone) + 1% penicillin/streptomycin (Gibco) and kept at 37°C, 5% CO_2_.

### METHOD DETAILS

#### Plasmids

The lentiviral destination vector pLX302 (Addgene #25896) was recombined with pDONR221 EGFP (Addgene #25899) or pDONR223 MX1, OAS1, OAS2, and OASL from the Human ORFeome v5.1 (Yang et al., 2011) to yield pLX302 EGFP-V5 puro (Addgene #158644), pLX302 MX1-V5 puro (Addgene #158640), pLX302 OAS1-V5 puro (Addgene #158641), pLX302 OAS2-V5 puro (Addgene #158642), and pLX302 OASL-V5 puro (Addgene #158643). pDONR223 MAVS Glu^93^Gln^148^Gln^271^ was obtained from the human ORFeome v5.1 (Yang et al., 2011), originally derived from IMAGE clone #5751684. MAVS Glu^93^Gln^148^Gln^271^ amplicon was prepared with SpeI and MfeI restriction sites and cloned into pEN_TTmiRc2 3xFLAG (Addgene #83274) that had been digested with SpeI and MfeI (Addgene #158628). pDONR223 MAVS Glu^93^Ala^148^Gln^271^ was obtained by site-directed mutagenesis of pDONR223 MAVS Glu^93^Gln^148^Gln^271^ (Yang et al., 2011). MAVS Glu^93^Ala^148^Gln^271^ amplicon was prepared with SpeI and MfeI restriction sites and cloned into pEN_TTmiRc2 3xFLAG (Addgene #83274) that had been digested with SpeI and MfeI (Addgene #158630). pEN_TT-3xFLAG-MAVS Glu^93^Gln^148^Ala^271^ (Addgene #158631) and pEN_TT-3xFLAG-MAVS Gln^93^Gln^148^Gln^271^ (Addgene #158629) were obtained by site-directed mutagenesis of the pEN_TT-3xFLAG-MAVS Glu^93^Gln^148^Gln^271^ plasmid (Addgene #158628). pEN_TT-3xFLAG-MAVS Glu^93^Ala^148^Ala^271^ (Addgene #158633) was obtained by site-directed mutagenesis of the pEN_TT-3xFLAG-MAVS Glu^93^Ala^148^Gln^271^ plasmid (Addgene #158630). pEN_TT-3xFLAG-MAVS Gln^93^Gln^148^Ala^271^ (Addgene #158632) was obtained by site-directed mutagenesis of the pEN_TT-3xFLAG-MAVS Glu^93^Gln^148^Ala^271^ plasmid (Addgene #158631). Site-directed mutagenesis of pEN_TT-3xFLAG-MAVS Glu^93^Gln^148^Ala^271^ consistently caused one of two off-target mutations. Two clones with different off-target mutations were digested with MfeI and NaeI. The fragments were gel purified, and the fragments with the non-off-target mutations were ligated together to yield pEN_TT-3xFLAG-MAVS Gln^93^Gln^148^Ala^271^ (Addgene #158632). All site-directed mutagenesis was performed with the QuikChange II XL kit (Agilent).

pEN_TT donor vectors were recombined into pSLIK hygro by LR recombination to obtain pSLIK 3xFLAG-MAVS Glu^93^Gln^148^Gln^271^ hygro (Addgene #158634), pSLIK 3xFLAG-MAVS Glu^93^Ala^148^Gln^271^ hygro (Addgene #158636), pSLIK 3xFLAG-MAVS Glu^93^Gln^148^Ala^271^ hygro (Addgene #158637), pSLIK 3xFLAG-MAVS Gln^93^Gln^148^Gln^271^ hygro (Addgene #158635), pSLIK 3xFLAG-MAVS Glu^93^Ala^148^Ala^271^ hygro (Addgene #158639), and pSLIK 3xFLAG-MAVS Gln^93^Gln^148^Ala^271^ hygro (Addgene #158638).

DNA for CVB3 was obtained by partial digestion of CVB3-M1 (Kandolf and Hofschneider, 1985) (kindly provided by K. Klingel) with EcoRI, gel purification of the full-length CVB3 insert, and ligation into pcDNA3 digested with EcoRI to obtain pcDNA3 CVB3 (Addgene 158645). The shRNA targeting SV40 T antigen (GCATAGAGTGTCTGCTATTAA) was custom designed through the Genetic Perturbation Platform web portal (https://portals.broadinstitute.org/gpp/public/). The oligos were modified to contain a PstI- containing loop, self-annealed, and ligated into pLKO.1 neo (Addgene #13425) to yield pLKO.1 shSV40 neo (Addgene #158646).

#### Viral Transduction

pSLIK MAVS alleles, pSLIK LacZ, pSLIK Luc, and pLKO.1 shSV40 were packaged as lentivirus in HEK293T cells by calcium phosphate precipitation with an EGFP-expressing plasmid as a positive control. AC16-CAR cells were transduced with lentivirus + 8 μg/ml polybrene in 6-well dishes and allowed to grow for 48 hours. Transduced pSLIK cells were then transferred to 10- cm dishes and selected with 10 μg/ml blasticidin (to maintain CAR-V5 expression) and 100 μg/ml hygromycin until control plates had cleared. Transduced shSV40 cells were used unselected.

#### AC16 Quiescence

For studies involving quiescent cells (Figures 2B, 2C, and S2B–D only), AC16 or AC16-CAR cells (70,000 per well) were plated on a 12-well dish precoated with 0.02% (w/v) gelatin for 1–2 hours at 37°C. After 24 hours, cells were washed with PBS and the culture medium was changed to differentiation medium: DMEM/F12 (Gibco) + 2% horse serum (Thermo Fisher) + 1x Insulin–Transferrin–Selenium supplement (Thermo Fisher) + 1% penicillin–streptomycin (Gibco). After 24 hours, cells were washed with PBS and transduced with shSV40 lentivirus (125 µl per well) + 8 µg/ml polybrene in a total volume of 500 µl. After 18–21 hours, cells were washed with PBS and refed with differentiation medium. After another 24 hours, cells were washed with PBS and refed with differentiation medium again, and cells were lysed 24 hours after the final refeed.

#### CVB3 Infection

AC16-CAR cells were plated at 50,000 cells/cm^2^ for 24 hours. AC16-CAR cells expressing inducible MAVS Glu^93^Gln^148^Gln^271^, Glu^93^Gln^148^Ala^271^, Glu^93^Ala^148^Ala^271^, or Gln^93^Gln^148^Gln^271^ or LacZ were plated at 25,000 cells/cm^2^ for 24 hours then treated with 1 μg/ml doxycycline for 24 hours. Before CVB3 infection, 75% of the cell culture medium was removed, and cells were infected with CVB3 at the indicated MOI for one hour. During the infection, the plates were incubated at 37°C and rocked every 10–15 minutes to ensure even coverage of the virus. After one hour, the media was aspirated, and cells were refed with fresh AC16 growth medium lacking selection antibiotics. At the end of the infection period, the conditioned media containing released virions was collected, centrifuged at 2,500 rcf to spin out dead cells and debris, and stored at –80°C for viral titering.

#### Cell Morphometry

Cell dimensions were analyzed by staining terminally CVB3-infected cells as before (Jensen et al., 2013). Briefly, AC16-CAR cells were infected with CVB3 at 10 MOI for 24 hours, stained with the LIVE/DEAD Fixable Violet Dead Cell Stain Kit (Thermo Fisher), quenched with 10 mM glycine in PBS for five minutes, solvent fixed–permeabilized with 100% ice-cold methanol, and counterstained with DRAQ5 (Cell Signaling Technology). Imaging at late times after CVB3 infection allowed VROs to fuse and grow above the diffraction limit for fluorescence imaging (Melia et al., 2019). One hundred Violet-positive cells were imaged with an Olympus BX51 fluorescence microscope with a 40x 1.3 numerical aperture UPlanFL oil immersion objective and an Orca R2 charge-coupled device camera (Hamamatsu) with 2×2 binning. Images were thresholded in ImageJ to identify Violet-positive membrane borders, Violet-negative VRO borders, and DRAQ5-positive nuclear borders. When calculating cellular dimensions, we assumed a height equal to the diameter of a sphere corresponding to the average aggregate VRO volume per cell.

#### Cell Lysis

For immunoblotting, cells were washed with ice-cold PBS and lysed in radioimmunoprecipitation buffer plus protease–phosphatase inhibitors: 50 mM Tris-HCl (pH 7.5), 150 mM NaCl, 1% (v/v) Triton X-100, 0.5% (w/v) sodium deoxycholate, 0.1% (w/v) SDS, 5 mM EDTA, 10 µg/ml aprotinin, 10 µg/ml leupeptin, 1 µg/ml pepstatin, 1 µg/ml microcystin-LR, 200 µM Na_3_VO_4_, and 1 mM PMSF. Protein concentrations of clarified extracts were determined with the BCA Protein Assay Kit (Pierce).

For RNA analysis, culture medium was aspirated and cells were immediately lysed in Buffer RLT Plus. Total RNA was purified with the RNEasy Mini Plus kit (Qiagen) as recommended. RNA concentrations were determined by absorption spectrophotometry on a Nanodrop.

#### Immunoblotting

Quantitative immunoblotting was performed on 10, 12, or 15% polyacrylamide gels with tank transfer to polyvinylidene difluoride membrane and multiplex near-infrared fluorescence detection as described previously (Janes, 2015). Primary antibodies were used at the following dilutions: VP1 (Mediagnost, 1:2000 dilution), eIF4G (Cell Signaling Technology, 1:2000 dilution), V5 epitope tag (Invitrogen, 1:5000 dilution or Bethyl, 1:5000 dilution), HSP90 (Santa Cruz Biotechnology, 1:2000 dilution), tubulin (Abcam, 1:20,000 dilution or Cell Signaling Technology, 1:2000 dilution), p38 (Santa Cruz Biotechnology, 1:5000 dilution), vinculin (Millipore, 1:10,000 dilution), IκBα (Cell Signaling Technology, 1:2000 dilution), phospho-TBK1 (Cell Signaling Technology, 1:1000 dilution), total TBK1 (Cell Signaling Technology #3504), phospho-IRF3 (Cell Signaling Technology, 1:1000 dilution), total IRF3 (Cell Signaling Technology, 1:1000 dilution), MDA5 (Abcam, 1:1000 dilution), MAVS (Cell Signaling Technology, 1:1000 dilution), actin (Ambion, 1:5000 dilution), FLAG (Sigma-Aldrich, 1:5000 dilution). Serial dilutions of V5-containing Multiple Tag (GenScript) were used for absolute quantification of CAR-V5.

#### In Vitro Transcription of CVB3 Positive and Negative Strands

The pcDNA3 CVB3 plasmid was linearized at the 3’ end with NotI (positive-strand template) or at the 5’ end with SnaBI (negative-strand template), gel purified, and used for in vitro transcription with T7 (positive strand) or SP6 (negative strand) RNA polymerase from the MAXIscript SP7/T7 kit (Ambion). In vitro transcription reactions were incubated at 37°C for two hours before the template was digested with TURBO DNAse I at 37°C for 15 minutes and transcripts purified by LiCl precipitation according to the manufacturer’s recommendations (Thermo Fisher). RNA concentrations were determined by absorption spectrophotometry on a Nanodrop.

#### dsRNA Transfection

Poly(I:C) (High Molecular Weight) and 5’ppp-dsRNA control were prepared according to the manufacturer’s separate recommendations (Invivogen). dsCVB3 was prepared by mixing equal volumes of CVB3 positive and negative strand at 100 ng/µl in 0.9% (w/v) NaCl, denaturing for at 68°C for 10 minutes, and cooling to 25°C at 0.1°C per second on a thermocycler.

For dsRNA transfection, AC16-CAR cells were plated at 50,000 cells/cm^2^ on a 12-well plate overnight and transfected with up to 1 µg dsRNA complexed with 3 µl Lipofectamine 3000 + no P3000 reagent (Invitrogen) in 200 µl total volume of DMEM/F12 (Gibco). For dose responses, lipocomplexes were serially diluted threefold in DMEM/F12 over ∼two decades before addition to AC16-CAR cells. Four hours after transfection, cells were lysed in 350 µl Buffer RLT Plus and purified with the RNEasy Plus Mini Kit (Qiagen). Purified RNA concentrations were determined by absorption spectrophotometry on a Nanodrop.

#### ISG Transfection and Protein-Synthesis Inhibition

To estimate ISG cellular half-lives, 293T/17 cells were plated at 100,000 cells/cm^2^ on a 24-well plate and transfected with 1 µl Lipofectamine 3000 + 1 µl P3000 reagent (Invitrogen) and 250 ng pLX302 EGFP-V5 puro + 250 ng pLX302 ISG-V5 puro (ISG = MX1, OAS1, OAS2, or OASL) in 100 µl total volume of DMEM (Gibco). Twenty-four hours after transfection, cells were treated with 50 µM cycloheximide (Sigma-Aldrich) for the indicated times and lysed in 50 µl RIPA + PPIs for immunoblot analysis of ∼30 µg extract for V5 epitope tag (Invitrogen, 1:5000 dilution [MX1-V5] or Bethyl, 1:5000 dilution [OAS1-V5, OAS2-V5, and OASL-V5]) and HSP90 (Santa Cruz Biotechnology, 1:2000 dilution)–tubulin (Cell Signaling Technology, 1:2000 dilution)–p38 (Santa Cruz Biotechnology, 1:5000 dilution) as loading controls. Cotransfection of pLX302 EGFP-V5 yielded a short-lived FP-V5 truncation that served as a positive control for cycloheximide efficacy (FP_control_).

#### RNA-seq and Analysis

500 ng total RNA was prepared using the TruSeq Stranded mRNA Library Preparation Kit (Illumina). Samples were sequenced on a NextSeq 500 instrument with NextSeq 500/550 Mid Output v2 kit (150 cycles; Illumina). Pooled samples were run in triplicate to obtain a minimum sequencing depth of 20 million reads per sample. Adaptors were trimmed using fastq-mcf in the EAutils package (version 1.1.2) with the following options: -q 10 -t 0.01 -k 0 (quality threshold 10, 0.01% occurrence frequency, and no nucleotide skew causing cycle removal). Datasets were aligned to the human (GRCh38) genome with additions (CVB3 genome and SV40 genome) using HISAT2 (version 1.2.0) with the following options: --dta (downstream transcriptome assembly for subsequent assembly step) and --rna-strandedness RF (for paired-end reads generated by the TruSeq strand–specific library). Output SAM files were converted to BAM files using samtools (version 1.4.1). Alignments were assembled into transcripts using StringTie (version 2.0.6) with the following options: -e (to restrict assembly to known transcripts in the provided annotation) and -B (to save additional files for Ballgown). Differential gene expression analysis was carried out using edgeR (version 3.28.1) on raw read counts that passed the abundance-filtering step. Abundance filtering was performed by the cpm function in edgeR to retain transcripts that were expressed at greater than 100 counts per million in at least one cell line. Trimmed mean of M values normalization using the calcNormFactors function before differential expression analysis using exactTest in edgeR. The 1952 transcripts that were commonly differentially expressed [5% false discovery rate (FDR)] between AC16 parental and AC16-CAR cells are shown in Figure 2B.

#### HeLa Cell Transcriptomic and Proteomic Data

Raw HeLa cell transcriptomic data (GSE111485) was downloaded from the GEO sequence read archive. The data were aligned and TPM calculated exactly as for the AC16 and AC16-CAR cells. “HeLa12” and “HeLa14” were profiled in triplicate, so each of their TPM values were averaged. Proteomic data was obtained from https://helaprot.shinyapps.io/crosslab/ by searching for CD55 for DAF and CXADR for CAR. The 14 HeLa samples were paired by numeric index with the 14 HeLa TPM samples obtained from GEO. Obtained proteomic data was in log_10_ format, so absolute protein abundances were calculated by 10^(log10_protein)^.

#### Determination of CVB3 Particle-to-PFU Ratio

Titered production stocks of CVB3 prepared in permissive HeLa cells were ultracentrifuged at 100,000 rcf for one hour at 4°C and lysed in Buffer RLT Plus for total RNA purification with the RNEasy Mini Plus kit (Qiagen) as recommended. RNA concentrations were determined by absorption spectrophotometry on a Nanodrop. Conditioned medium from uninfected cells was used as a negative control to confirm specificity to viral RNA. Similar results were obtained when CVB3 was ultracentrifuged at 120,000 rcf for 18 hours at 8°C through a 30% sucrose cushion and into a glycerol button (Kim et al., 2005).

#### Quantitative PCR (qPCR)

Purified total RNA was digested with TURBO DNAse (Thermo Fisher) as recommended, and 500 ng of DNAse-treated RNA was reverse transcribed with 2.5 µM oligo(dT)_24_ primer as described previously (Miller-Jensen et al., 2007). *PRDX6*, *HINT1*, and *GUSB* were used as loading controls, and *GAPDH* was used as a fourth housekeeping gene to confirm accuracy of the loading normalization.

#### Tagged Strand-Specific qPCR

Purified total RNA was digested with TURBO DNAse (Thermo Fisher) as recommended, and 250 ng of DNAse-treated RNA was reverse transcribed with 1 pmol biotinylated strand-specific primer (positive strand: 5’biotin-GGGTGTTCTTTGGATCCTTG; negative strand: 5’biotin-TGCAACTCCCATCACCTGTA), 500 µM dNTPs, 5 mM DTT, 20 U RNAsin Plus RNAse inhibitor (Promega), and 100 U Superscript III reverse transcriptase (Invitrogen) in 1x first-strand buffer as recommended in a total volume of 10 µl. After reverse transcription for 60 minutes at 55°C, samples were heat inactivated for 15 minutes at 70°C and RNA was digested with 2.5 U RNAse H (NEB) for 20 minutes at 37°C, followed by heat inactivation for 20 minutes at 65°C. Purified standards—either RNA from CVB3 virions purified by ultracentrifugation (positive strand) or in vitro transcribed RNA from SnaBI-digested pcDNA3 CVB3 (negative strand)—were added in the range of 10^8^–10^3^ copies per 10 µl reverse transcription along with 250 ng of DNAse-treated RNA from cells that had not been infected with CVB3.

Biotinylated cDNA was purified with 5 µl streptavidin magnetic beads (Thermo Fisher) that had been washed twice with 1x first-strand buffer and resuspended to the original storage volume. Streptavidin beads were incubated with the first-strand reaction for one hour with tapping every 15 minutes. The 15 µl samples were diluted with 40 µl PCR-grade water, magnetized for one minute, and then washed three times with 50 µl PCR-grade water. After the final wash, the beads were resuspended in 50 µl and then 1/10th of the nonbead volume (4.5 µl) was measured by qPCR with 2.5 pmol each of CVB3-specific primers (Table S2) as described previously (Miller-Jensen et al., 2007).

#### Plaque Assay

Conditioned medium from CVB3-infected AC16-CAR cells was collected and diluted 5 x 10^4^ or 5 x 10^5^ in serum-free DMEM depending on the titer. A confluent 6-well plate of permissive HeLa cells was washed twice with serum-free DMEM, then 200 µl of diluted conditioned medium or serum-free DMEM was added to the cells. The cells were incubated at 37°C for one hour with rocking every 10–15 minutes to ensure even coverage of the virus. After one hour, the medium was aspirated and cells were washed twice with serum-free DMEM. Sterile 1.5% (w/v) agar in water was mixed with 2x DMEM in a 1:1 ratio and added to the wells to form an agar plug. The cells were incubated at room temperature 10–15 minutes until solidification of the agar plug.

Cells were incubated at 37°C for 48–65 hours depending on the clearance rate of the plaques. Once plaques reached a countable size, 2 ml of 2% (w/v) formaldehyde was added to each well for 15 minutes to fix the cells. After fixation, the agar plugs were carefully pulled out using a plastic spatula. Fixed cells were stained with 0.5% (w/v) crystal violet in 25% (v/v) methanol in Milli-Q water for 10 minutes. Crystal violet staining was stopped by submerging the plates three times into a 1% bleach solution, and then wells were gently rinsed with Milli-Q water three times from a squirt bottle. The plates were air dried and scanned on a Licor Odyssey scanner in the 700 channel.

#### 3Cpro Sequence Logo

We recreated the sequence logo for enteroviral 3C^pro^ substrates by assembling the 202 3C^pro^ cleavage sites from 27 SwissProt accession numbers in Table 6 of the original publication describing NetPicoRNA (Blom et al., 1996). Flanking sequences were amended with the latest revisions in UniProt, and the vectorized sequence logo was generated with WebLogo (Crooks et al., 2004).

#### CVB3 Complete Kinetic Model

The complete kinetics of CVB3 infection were described by 54 coupled differential equations that were organized into modules for delivery, replication, and encapsidation. After construction, the modules were interconnected by viral inputs–outputs and host-cell feedback. The system of differential equations was solved numerically with the ode15s function in MATLAB. The two core functions are 1) CVB3ODEEval.m, which defines the initial conditions and rate parameters, takes the user-defined inputs (CVB3 dose, simulation parameters, output options), and performs the bookkeeping on viral species (RNA strands, replicative intermediates, and protein classes); and 2) CVB3ODEfunc.m, which defines the rate equations, the switch to VROs, and the antiviral sensing-transduction mechanism. Additional functions provide capabilities for systematic plotting and sensitivity analysis (Data and Code Availability). MATLAB version 2019b or later is required to handle the export of tabular data, and the Statistics Toolbox is required to display sensitivity analyses. The following Method Details explain and justify the key assumptions within each module, feedback, and bookmarked species.

##### Delivery Module

At the start of a single-cell infection, a user-defined number of PFUs was placed in the restricted cell-surface volume defined by the estimated cell surface area multiplied by the 30-nm diameter of a CVB3 capsid (Table S1). For a cell-population infection, cell-to-cell variation in response to a viral inoculum follows a Poisson distribution (Ellis and Delbruck, 1939). Therefore, a Poisson-distributed pseudorandom number (representing a single-cell PFU) was drawn at each simulation with *λ* equal to the user-defined MOI. Cellular DAF and CAR were also concentrated in the restricted cell-surface volume for CVB3 binding and trafficking. Clustering of DAF was embedded in the rate of DAF trafficking to tight junctions, which used a kinetic parameter drawn from observations that relocalization is complete within 25–30 minutes (Coyne and Bergelson, 2006). Tight junctions were not encoded as a volume-restricted compartment but rather as their own set of species on the plasma membrane. DAF-bound CVB3 in tight junctions (*bDAF_TJ_*) was allowed to unbind-rebind in the tight junction or transfer CVB3 directly to CAR:

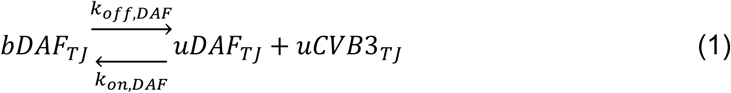

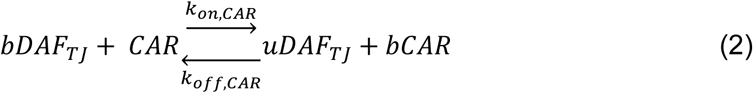

In addition, CAR could bind CVB3 that had dissociated in tight junctions (*uCVB3_TJ_*):

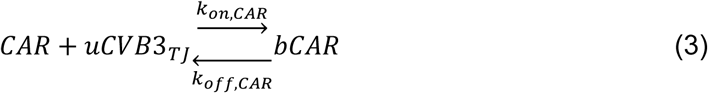

The forward rates for Equations 2 and 3 were assumed to be identical to place equal weight on the two possible paths to CAR-bound CVB3. CAR internalizes with CVB3 (Chung et al., 2005) and we assumed that it was not recycled; however, this assumption was inconsequential to any of the kinetics (Figure S2A).

Entry of CVB3 occurs more slowly than poliovirus (Brandenburg et al., 2007; Chung et al., 2005). Therefore, delivery of CVB3 to the cytoplasm was encoded as two first-order processes in series, the first using a standard endocytosis rate and the second capturing pore formation and release of viral RNA (Table S1). Defective viral particles were assumed to bind to DAF–CAR and deliver to the cytoplasm identically to infectious PFUs.

##### Replication Module

Mammalian cells contain 10^6^–10^7^ ribosomes (Milo et al., 2010). Considering competition with newly transcribed host-cell RNAs and with RNA from defective particles, we assumed that 10^5^ ribosomes per cell were theoretically accessible to the infectious virus. This number is also roughly equivalent to the number of eIF4G molecules [∼10^5.5^ copies per HeLa cell (Liu et al., 2019), with TPM values comparable in AC-16 cells], allowing us to equate eIF4G–ribosome as a single species in the model. We further simplified by considering viral translation in terms of polysomes containing *N* ribosomes. Each polysome translated viral polyprotein *N* times more rapidly but formed and released a translational complex with viral RNA *N* times more slowly than an individual ribosome. Translational initiation was emphasized as a rate-limiting step by selecting an operational early-polysome size of *N* = 2.5 (Figure S3A). Defective viral RNAs in the cytoplasm were assumed not to interact with the ribosome pool that was theoretically accessible, but this assumption was not critical to infection outcomes (Figures S3E and S3F).

Among these polysomes, 80–90% would be actively translating host-cell RNAs and therefore be inaccessible at the time of infection (Milo et al., 2010). The conversion of polysomes from inaccessible to accessible was encoded as an enzyme-catalyzed transition rate dependent on the concentration of viral protease, the concentration of inaccessible ribosomes (i.e., uncleaved eIF4G), and Michaelis-Menten parameters for rhinoviral 2A^pro^ (Wang et al., 1998) (Table S1). The reported K_M_ for enterovirus 71 3C^pro^ is equivalently far above endogenous substrates and shows comparable catalytic efficiency (Shang et al., 2015), enabling 2A^pro^–3C^pro^ to be lumped as a single viral protease.

The dissociation of a translation complex is non-intuitive and thus explained in detail here. The rate of protein production used poliovirus translation rates per amino acid multiplied by the length of the CVB3 polyprotein (Dorner et al., 1984). Each translation event yields one 3D^pol^ bound to VRO, one lumped protease, ⅕ of a capsid pentamer at the VRO (which couples to ⅕ of a 2C^ATPase^), and ⅘ of unbound 2C^ATPase^ at the VRO to balance polyprotein stoichiometry. At a rate *N*^-1^ times the translation rate, the translation complex dissociates to release a positive-strand RNA at the VRO (Goodfellow et al., 2000; Novak and Kirkegaard, 1994) and repopulate the available polysome pool by *N* ribosomes. Placing a translation-competent positive strand at the VRO mimics cis replication (Goodfellow et al., 2000; Novak and Kirkegaard, 1994), in that defective viral RNAs do not reach this step in the complete kinetic model. More-direct alternatives, such as coupling 3D^pol^ with released positive-strand RNA as a replicative intermediate at the VRO, gave rise to premature increases in negative strand that were not observed by experiment (Figure 3C).

The early steps of VRO formation remain elusive and were specified heuristically in the complete kinetic model. We assumed a switch-like transition from solution-phase behavior in the cytoplasm to the restricted surface volume of the VRO defined by the estimated VRO surface area multiplied by the 7.1-nm height of a 3D^pol^ molecule (Table S1). Based on these calculations, the concentrating effect (*C_E_*) of shuttling a molecule from the cytoplasm to the VRO surface volume was 3216-fold. Degradation rates at the VRO surface were reciprocally scaled down by *C_E_* to yield the same turnover as in the cytoplasm. The switch-like transition to VROs at 25 molecules of 3D^pol^ created a transient stoichiometric imbalance at the start of the transition (Figure S4B). The shift occurred when the rate processes of VRO-resident species were instantly scaled up and their whole-cell concentrations were reciprocally scaled down. The imbalances were short-lived, and alternate encodings of the VRO transition, such as a ramped transition, did not satisfactorily capture the measured protein–RNA dynamics.

At the VRO, viral RNA interacted with 3D^pol^ at the slow rate of association measured for poliovirus 3D^pol^ (Arnold and Cameron, 2000). The rate of RNA-dependent RNA polymerase elongation was likewise drawn from poliovirus (Castro et al., 2007). We did not postulate any differences in association or elongation between positive- and negative-strand templates (Regoes et al., 2005). However, we assumed that negative strand was not released from a replicative intermediate after replication of the positive strand was complete and had been released. This assumption reflects the cooperative RNA binding that results from 3D^pol^ oligomeric arrays on the VRO surface (Lyle et al., 2002). Positive strands, by contrast, must be released for encapsidation.

##### Encapsidation Module

The encapsidation module leveraged the work of Zlotnick and colleagues for modeling viral capsid assembly (Endres and Zlotnick, 2002; Li et al., 2012; Zlotnick, 2003). During assembly, contact affinities that are weak individually (∼1 mM) build upon one another through multivalency among symmetric subunits with multiple binding sites (Zlotnick, 2003). Using established formalisms (Endres and Zlotnick, 2002), we calculated effective pseudocritical concentrations (apparent dissociation constants) for all realistic combinations of pentamer and contact sites in a 12-pentamer virus assembly. We took the median effective pseudocritical concentration (∼1 µM, calculated with Capsid_assembly_script.m) as a representative affinity for any step during the assembly process from 1–12 pentamers. By the same analysis, a slight increase in the individual contact affinity (0.1 mM; Figure 4E) increased the median affinity to 50 nM (simulated by decreasing the off rate; Figure 4E). The simplification avoids the need to assume or inventory specific geometric paths to viral assembly while recognizing the typical binding energies involved.

Recruitment and retention of pentamers to the VRO occurred through a direct interaction of 1:1 stoichiometry with 2C^ATPase^ (Liu et al., 2010). Kinetic parameters for this interaction are not known, but the complete kinetic model was not very sensitive to a biologically plausible range of values (Figure 4F). Crucial to the stoichiometric balance was to keep track of each molecule of 2C^ATPase^ “consumed” by pentamer binding to the VRO and then “regenerated” when pentamers self-assembled. For example:

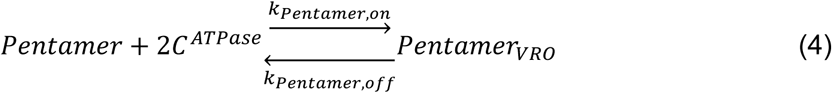

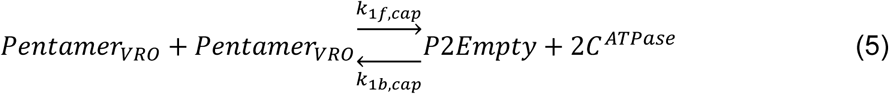

Pentamer assemblies were assumed to grow and shrink linearly, with individual pentamer or RNA–pentamer complexes recruited to or removed from an intermediate assembly.

The mechanisms retaining viral RNA in the VRO are heterogeneous, requiring a more- generic exchange rate between cytoplasmic and VRO compartments. The on and off rates were equal (*k_on_* = *k_off_* = 1 hr^-1^), but at the VRO, *C_E_* (= 3216) provided a driving force for RNA to leave that was delayed by the retention time implied by the rate of exchange:

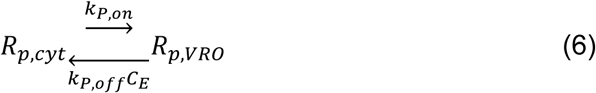

At these exchange rates, a typical RNA-protein association rate of 25 nM^-1^hr^-1^ (Gleitsman et al., 2017) yielded an effective membrane affinity of (3216 x 1 hr^-1^)/25 nM^-1^hr^-1^ ∼ 125 nM.

RNA and pentamer interacted at the VRO with the same kinetic parameters as the median pentamer–pentamer assemblies described above. We reduced combinatorics by assuming that RNA filling–unfilling of an intermediate assembly only occurred through gain or loss of an RNA–pentamer complex, which was defined to exist only at the VRO surface. For example:

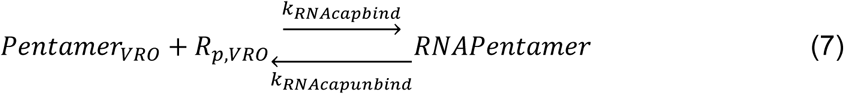

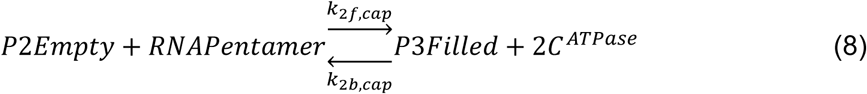

(Note the regenerated 2C^ATPase^ in Equation 8, consistent with the stoichiometric balance described in Equation 5.) The assembly of 12 pentamers with positive-strand RNA was considered the irreversible endpoint of encapsidation, which could be reached two ways:

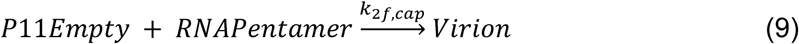

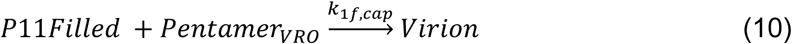

Empty provirions were assumed to be reversible:

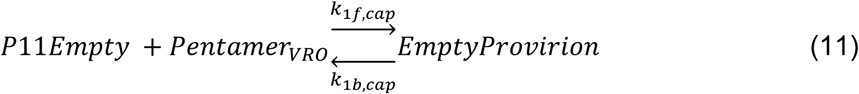

like the other steps in the encapsidation module. Rate parameters involving *Pentamer_VRO_* (*k_1f,cap_* and *k_1b,cap_*) were assumed to be identical to those involving RNAPentamer (*k_2f,cap_* and *k_2b,cap_*).

##### Antiviral and Antagonistic Feedbacks

The feedbacks overlaid on the CVB3 life cycle reflected known antiviral and antagonistic mechanisms, but they were not mechanistically encoded. Rather than striving for exact parameter values, we focused on lumped parameters and relationships that appropriately reflected dependencies and input–output characteristics. Future revisions could include more-elaborate detail when warranted by specific applications.

For the dsRNA sensor–transducer feedback leading to an interferon response, we defined a virus detection sigmoid with parameters (*K_D_, n_H_*) gleaned from the *IFNB* induction of cells lipofected with dsCVB3 (Figure 1B). To capture viral antagonism, the sigmoid maximum was scaled down hyperbolically depending on the concentration of viral protease and a half-effective concentration (*EC50_detectordeg_*) defined by qualitative system properties (Figure 5I):

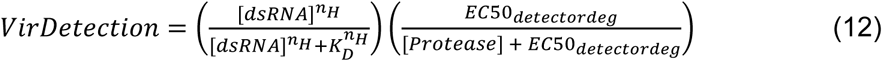

where [*dsRNA*] is the total concentration of replicative intermediates (Bookkeeping). This sigmoid was used to scale a maximum induced response of ISGs, which was defined by the maximum transcription rate for RNA polymerase II through a short ISG, multiplied by the current number of virus-inaccessible ribosomes (Table S1). For simulations involving supplemental interferon, *VirDetection* was superseded by *StimISG* = 1 (maximum stimulated interferon response) at the time of supplementation and maintained for the rest of the simulation.

The ISG response impinges on three facets of the viral life cycle. For i) the inhibition of viral translation, we hyperbolically down-scaled the formation rate of translation complexes:

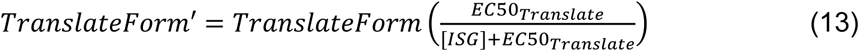

Although PKR (*EIF2AK2*) is highly abundant (∼480 TPM) and does not show interferon inducibility in AC16-CAR cells (Figure S1A), PKR is directly activated by dsRNA. Thus, the feedback architecture was assumed to be functionally similar to other ISGs. For ii) the deactivation of viral proteinases, we used a hyperbolic down-scaling function (similar to that of Equation 13) on the rate of viral protease production, assuming that protease deactivation was irreversible. For iii) the oligoadenylate synthetase–RNAse L acceleration of RNA turnover, we posited a maximum fold increase in the rate of turnover (*OAS_RNAdeg_* = 5) and scaled the basal rate of RNA degradation (*k_RNAdeg_*) hyperbolically to this maximum based on a half-effective concentration (*EC50_RNAdeg_*) and the concentration of ISGs:

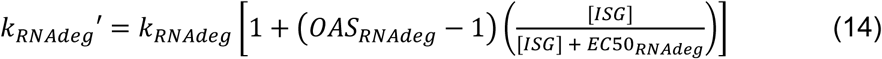

All feedback parameters were sampled lognormally about their best estimate with a user-defined log coefficient of variation as with the parameters and initial conditions of the complete kinetic model.

##### Bookkeeping

Total dsRNA concentrations combined the replicative intermediates using positive- or negative-strand templates. For ratiometric calculations where dsRNAs were counted with different efficiencies (Figure 3E), only the template strand was included in the calculation. Total viral protein included capsid from infectious PFUs and defective particles from internalization and all steps downstream (Figure 2A). Viral particles bound to DAF or CAR on the cell surface were assumed to be removed by the washing steps associated with cell lysis and protein analysis. Endosomal release of *R_p_* implied instant degradation of all associated capsid proteins. For RNA analysis, cells were not washed before lysis according to the manufacturer’s recommendation; therefore, these calculations included all viral particles bound to CAR on the cell surface (DAF affinity is so low that it was assumed to be removed by aspiration).

##### Sensitivity Analysis

Single-parameter sensitivity analysis was performed with SensitivityAnalysisfunc.m (Data and Code Availability) to alter individual model parameters over a 2^2^-fold range in either direction from the base value and record virion production at 24 hours.

#### MAVS Filamentation Model

The filamentation model for MAVS was described by 4–6 differential equations depending on the MAVS genotype:

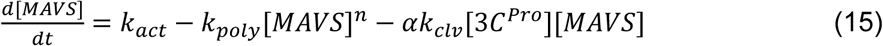

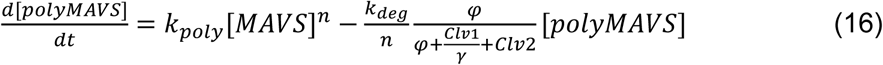

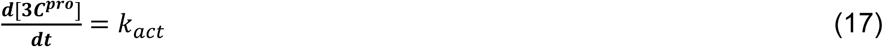

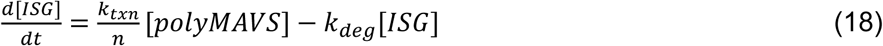

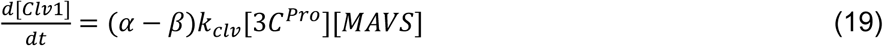

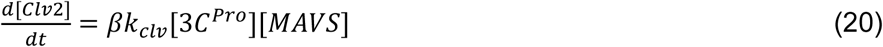

where *α* = 0, *β* = 0 for Glu^93^Ala^271^, *α* = 1, *β* = 0 for Glu^93^Gln^271^, and *α* = 2, *β* = 1 for Gln^93^Gln^271^. For the Gln^93^Gln^271^/Glu^93^Gln^271^ heterozygous model (Figure S7D), *α* = 1.5, *β* = 0.5, and for the Gln^93^Gln^271^ overexpression model in a Gln^93^Gln^271^/Glu^93^Gln^271^ heterozygous background (Figure S7C), *α* was set to 1 + 12.5/13 and *β* to 12.5/13 to reflect the ∼12-fold ectopic expression measured in Figure 6C.

Initial conditions were all set to zero, and rate parameters were nominally assumed to be fast (*k_poly_* = 1000), average (*k_act_*, *k_txn_* = 1), or slow (*k_clv_*, *k_deg_* = 0.1). *Φ* and *γ* were set to one as equal default feedback strengths, and *n* = 800 based on calculations from the literature (Wu et al., 2013; Wu et al., 2014). Filament assembly was assumed to occur by a two-step process used for other biological filaments (Lopez et al., 2016). Dynamic trajectories were solved numerically for 250 nominal time units with the ode15s function in MATLAB. Single-parameter sensitivities were evaluated by changing parameters individually 2^2^- or 10^2^-fold in either direction about the nominal parameter and using time-integrated ISG abundance as the model readout.

### QUANTIFICATION AND STATISTICAL ANALYSIS

#### dsRNA-Induced Interferon Response

Dose-independent induction of *IFNB*, *MX1*, and *OAS1* as a group (or *PKR* and *GAPDH* as a control group) was assessed by log-transformed three-way ANOVA of ISG transcripts with the following factors: dsRNA dose, transcript, and condition (control vs. poly(I:C) or control vs. dsCVB3). ANOVA *p* values testing a condition effect were Šidák corrected for multiple-hypothesis testing.

#### ISG Protein Half-Life Determination

Immunoblot bands were quantified by image densitometry as described previously (Janes, 2015). V5 band intensities were normalized to the averaged proportional loadings of HSP90, tubulin, and p38 for each sample on the immunoblot. Loading-normalized biological duplicates were averaged, and the ISG and FP_control_ time courses were normalized again by their respective maxima. Half-lives (*t_1/2_*) were estimated by nonlinear least-squares curve fitting in Igor Pro (WaveMetrics) to the following:

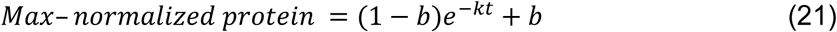

Where *b* represents a minimum baseline (estimated from the FP_control_ of each experiment and kept constant for the associated ISG) and *t_1/2_* = ln(2)/*k*.

#### Plaque Assay Calculations

Plaques were counted manually in ImageJ after contrast enhancement. The plaque forming units per milliliter (PFU/ml) was calculated from the following formula:

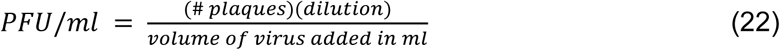

The dilution was either 5 x 10^4^ or 5 x 10^5^, and the volume of virus added to the wells was 0.2 ml.

#### Relative Cleavage of MAVS Genotypes

MAVS cleavage was assessed via densitometry in ImageJ as

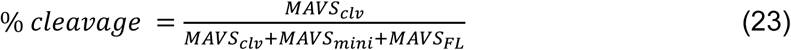

and normalized to MAVS Glu^93^Gln^148^Gln^271^ such that MAVS Glu^93^Gln^148^Gln^271^ cleavage = 1. Differences in cleavage were assessed by two-way ANOVA with CVB3 batch and genotype as factors.

#### Protein Abundance Estimation From TPM Data

Paired transcriptomic-proteomic data in HeLa cells (Liu et al., 2019) were related to one another using the following hyperbolic-to-linear equation:

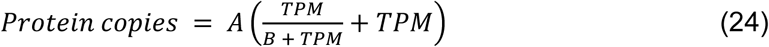

where *A* and *B* are fitted parameters. Equation 24 specifies a linear relationship between RNA and protein (Edfors et al., 2016) but allows for some nonlinearity at low RNA copy numbers.

Nonlinear least-squares parameterization was performed with the fminsearchbnd function in MATLAB. Asymptotic error analysis was used to calculate the 99% confidence interval of the fit. DAF and CAR protein copies were estimated in AC16 and AC16-CAR cells using the best-fit from the HeLa data for DAF or CAR, respectively.

## SUPPLEMENTAL INFORMATION TITLES AND LEGENDS

**Figure S1.**
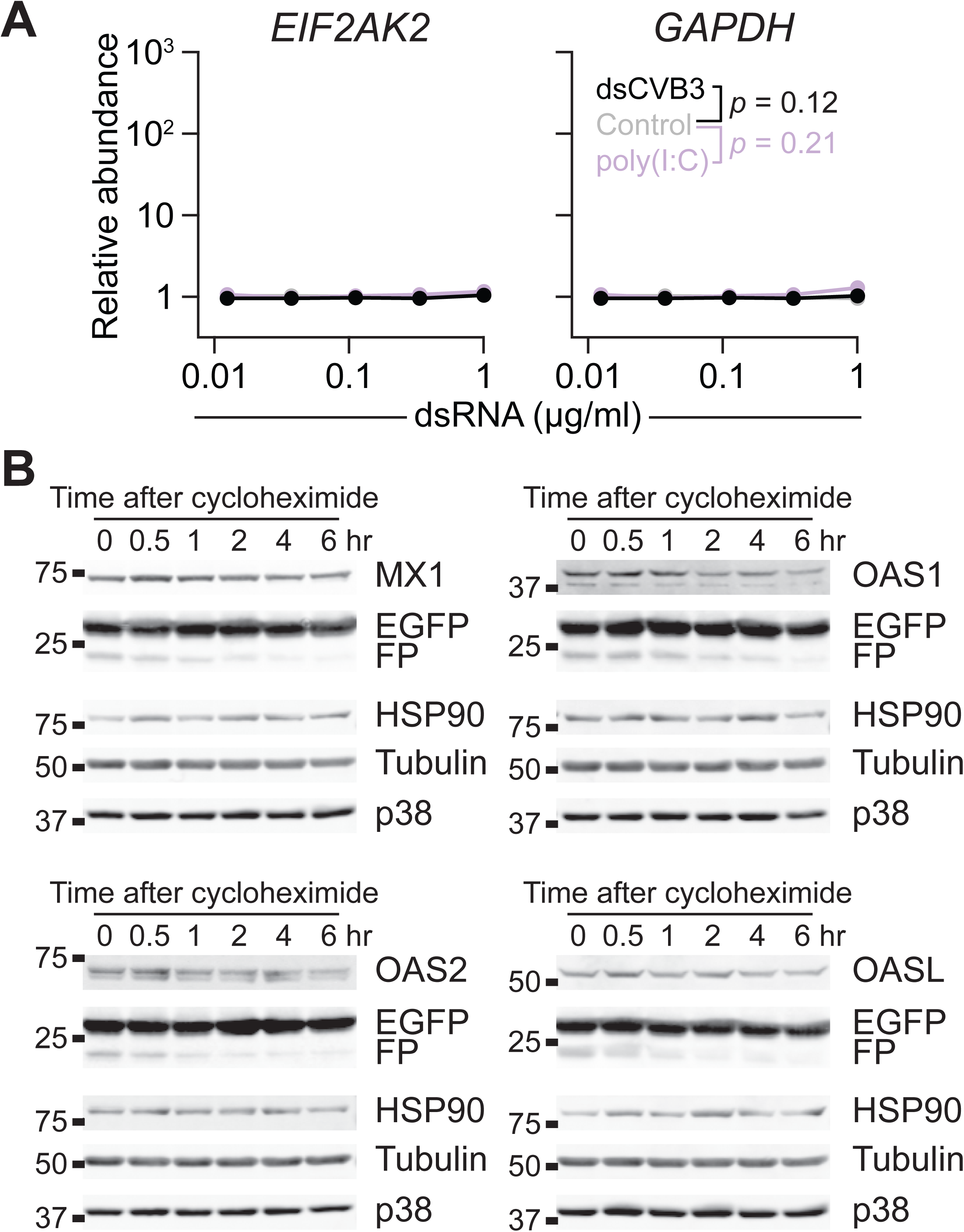
Related to Figure 1. Control genes for dsRNA induction and representative immunoblots of the cycloheximide experiment quantified in Figure 1C. (A) dsRNA lipofection does not give rise to nonspecific gene induction. AC16-CAR cells were lipofected with *n* = 5 doses of the indicated dsRNAs for four hours and analyzed for the indicated genes with *PRDX6*, *HINT1*, and *GUSB* used for loading normalization. Differences between conditions were assessed by Šidák-corrected, log-transformed three-way ANOVA. (B) Representative immunoblots used for half-life quantification. 293T/17 cells were lipofected with V5 epitope-tagged plasmids (equal mixtures of ISG and EGFP), treated with 50 µM cycloheximide for the indicated times, and lysed for quantitative immunoblotting. A C-terminal EGFP fragment (FP) was used as a control for a fast-degrading protein.

**Figure S2.**
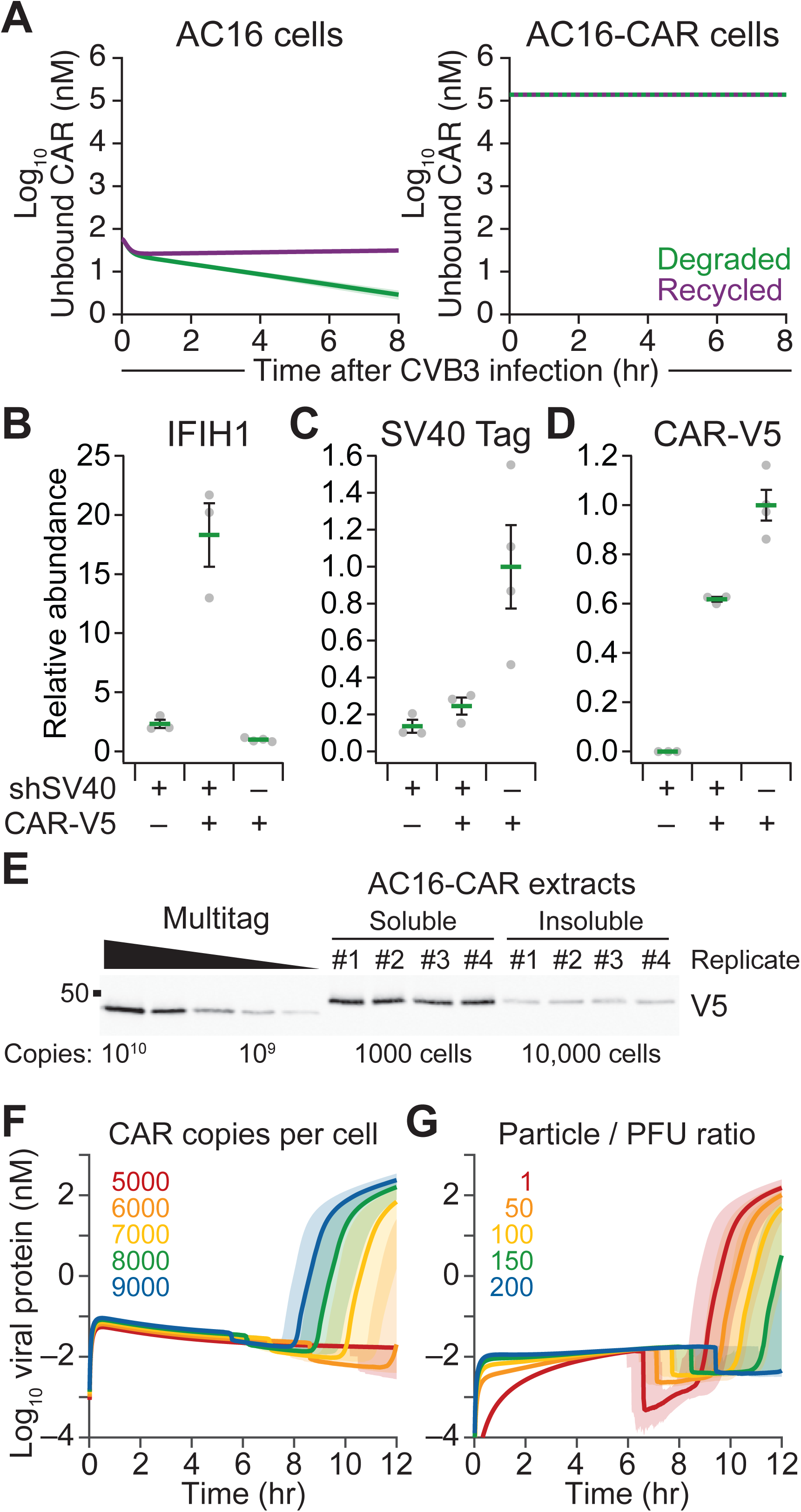
Related to Figure 2. Delivery controls, immunoblot quantification, V5 immunoblot image, and evaluation of receptor–ratio thresholds. (A) Recycling of CAR upon CVB3 internalization is inconsequential for permissive hosts. Nonpermissive (AC16, left) and permissive (AC16-CAR, right) complete kinetic models are simulated assuming total degradation (green) or recycling (purple) of CAR upon CVB3 internalization. AC16 cells are nonpermissive under both assumptions. Predictions of unbound CAR are shown as the median simulation ± 90% nonparametric confidence interval from *n* = 100 simulations of single-cell infections at 10 PFU with a parameter coefficient of variation of 5%. (B–D) Replicated densitometry of IFIH1, SV40 Tag, and CAR-V5. Data are shown as the mean ± s.e.m. from *n* = 3–4 biological replicates. (E) V5 immunoblot image of V5-containing Multitag recombinant protein along with RIPA-soluble (left) and RIPA-insoluble (right) extracts of AC16-CAR cells at the indicated cell abundance per lane. The abundance of soluble and insoluble CAR-V5 was combined per biological replicate in Figure 2F. (F) Minimum CAR abundance for productive infection of AC16 cells with CVB3. Predictions of total viral protein for 5000–9000 CAR molecules per cell are shown as the median simulation ± 90% nonparametric confidence interval from *n* = 100 simulations of single-cell infections at 10 PFU with a parameter coefficient of variation of 5%. (G) Critical particle-to-PFU ratio for accurately predicting the lack of CVB3 tropism in AC16 cells. Predictions of total viral protein for particle-to-PFU rations from 1–200 are shown as the median simulation ± 90% nonparametric confidence interval from *n* = 100 simulations of single-cell infections at 10 PFU with a parameter coefficient of variation of 5%.

**Figure S3.**
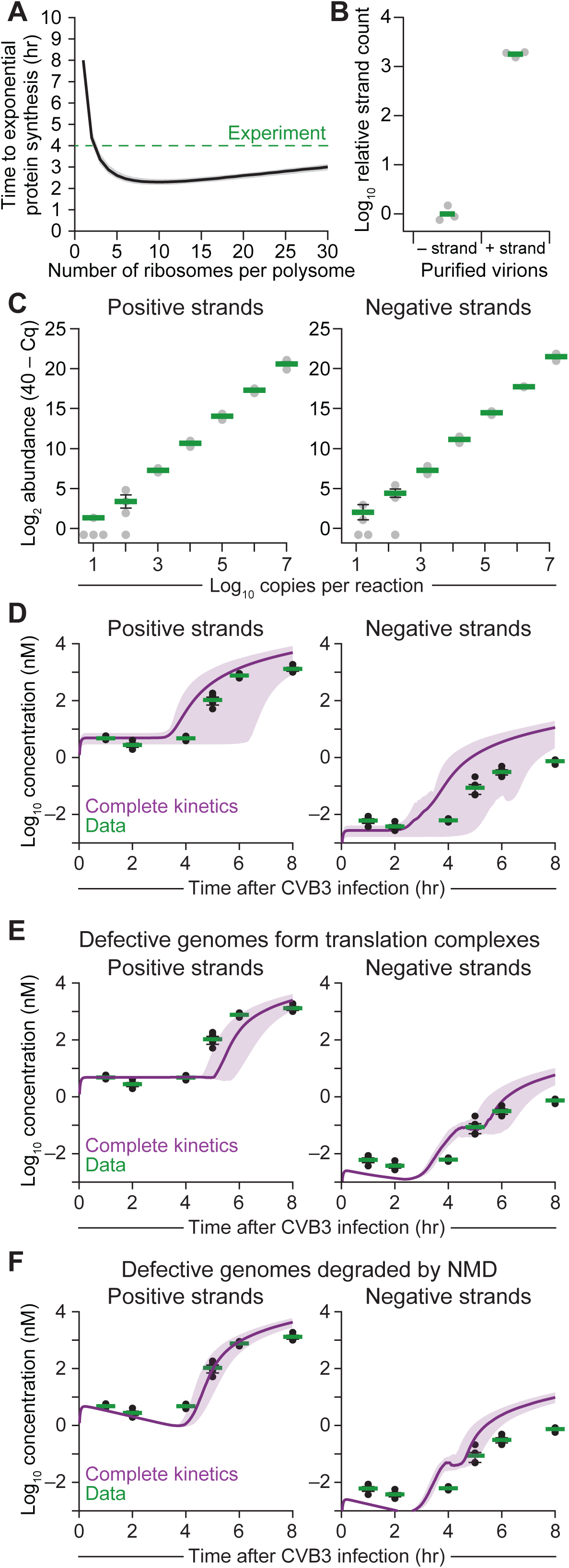
Related to Figure 3. Polysome size requirements, strand-specific tagged qPCR validation, and population-level simulations of viral RNA strand dynamics with complete kinetics. (A) Polysome size dictates the timing of viral protein synthesis. The start of exponential protein synthesis is shown as the median simulation ± 90% nonparametric confidence interval from *n* = 100 simulations of single-cell infections at 10 PFU with a parameter coefficient of variation of 5%. The experimentally observed timing (Figure 1D) is shown for reference (green). (B) Strand-specific tagged qPCR assessment of positive (+) and negative (–) strands in purified CVB3 virions. The measured strand ratio was 1790 ± 130. Data are shown as the geometric mean of *n* = 3 separate CVB3 dilutions of purified virion. (C) Quantitative precision and accuracy of the strand-specific tagged qPCR assay with positive strands from purified CVB3 virions and negative strands prepared by in vitro transcription. Data are shown as the mean log2 relative abundance [40 – qPCR quantification cycle (Cq)] (Singh et al., 2019) ± s.e.m. of assay quadruplicates at *n* = 7 separate positive-negative strand dilutions. Note the sporadic detection of the assay as the input approaches the single-molecule limit (assuming ∼10% conversion efficiency from RNA to amplifiable cDNA). (D) Complete kinetics captures the absolute viral RNA dynamics of positive and negative strands at the population level. (E and F) Alternative fates of defective CVB3 genomes do not substantially affect the modeled viral RNA dynamics of positive and negative strands. Defective genomes were assumed to (E) form defective polysome translation complexes that bound–unbound but did not yield polyprotein or (F) form defective polysome translation complexes that bound and instantly underwent nonsense-mediated decay (NMD). For (D–F), predictions (purple) were compared to data (green) obtained by strand-specific tagged quantitative PCR (qPCR) with purified standards. Data are reprinted from Figure 3C and shown as the geometric mean ± log-transformed standard error of *n* = 4 biological replicates of AC16-CAR cells infected at MOI = 10 for the indicated times. Predictions are shown as the mean simulation ± 90% nonparametric confidence interval from *n* = 100 simulations of population-level infections at MOI = 10 (D) or 100 simulations of single-cell infections at 10 PFU (E and F) with a parameter coefficient of variation of 5%.

**Figure S4.**
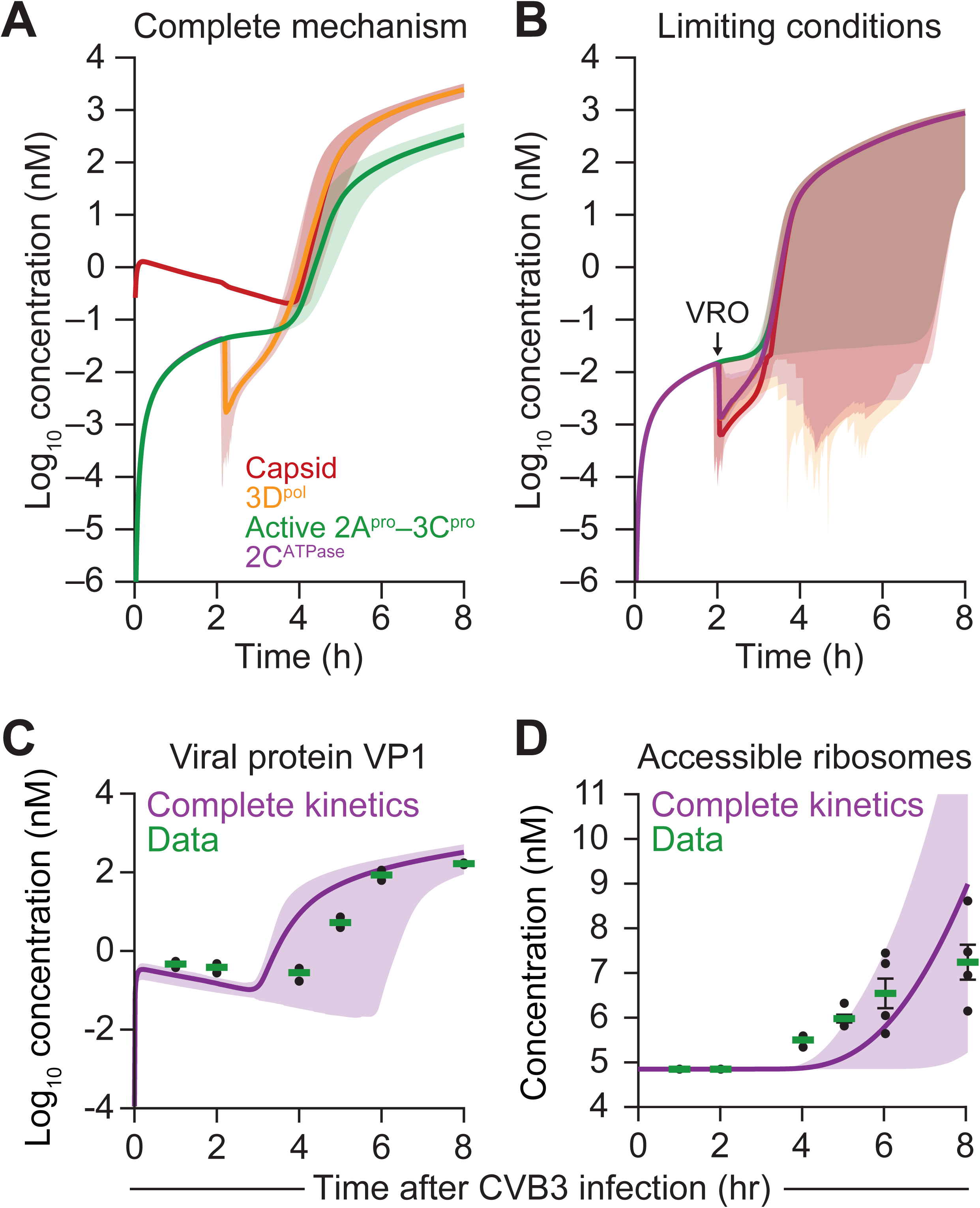
Related for Figure 4. Stoichiometric balance of CVB3 protein classes and population-level simulations of viral-protein and eIF4G cleavage dynamics. (A) Dynamics for the four protein species—capsid protein, 3D^pol^, active protease (2A^pro^–3C^pro^), and 2C^ATPase^—in the complete kinetic model. (B) Dynamics for the four protein species under the following limiting conditions for stoichiometric balance: no protein-RNA degradation, no antiviral response, no viral antagonism, no defective particles, and no bookkeeping of viral capsid from added PFUs. The separation of the four species at ∼two hours is caused by the immediate onset of the viral replication organelle (VRO) regime, which affects the apparent concentration of all species except active protease (STAR Methods). The solutions converge again within two hours. (C and D) Complete kinetics captures the dynamics of viral protein VP1 expression and accessible ribosomes (estimated from viral protease cleavage of host eIF4G) at the population level. Predictions (purple) were compared to data (green) obtained by quantitative immunoblotting. Data are reprinted from Figures 4B and 4C and shown as the geometric mean ± log-transformed standard error of *n* = 4 biological replicates of AC16-CAR cells infected at MOI = 10 for the indicated times. Predictions are shown as the mean simulation ± 90% nonparametric confidence interval from *n* = 100 simulations of population-level infections at MOI = 10 with a parameter coefficient of variation of 5%. For (A) and (B), predictions are shown as the median simulation ± 90% nonparametric confidence interval from *n* = 100 simulations of single-cell infections at 10 PFU with a parameter coefficient of variation of 5%.

**Figure S5.**
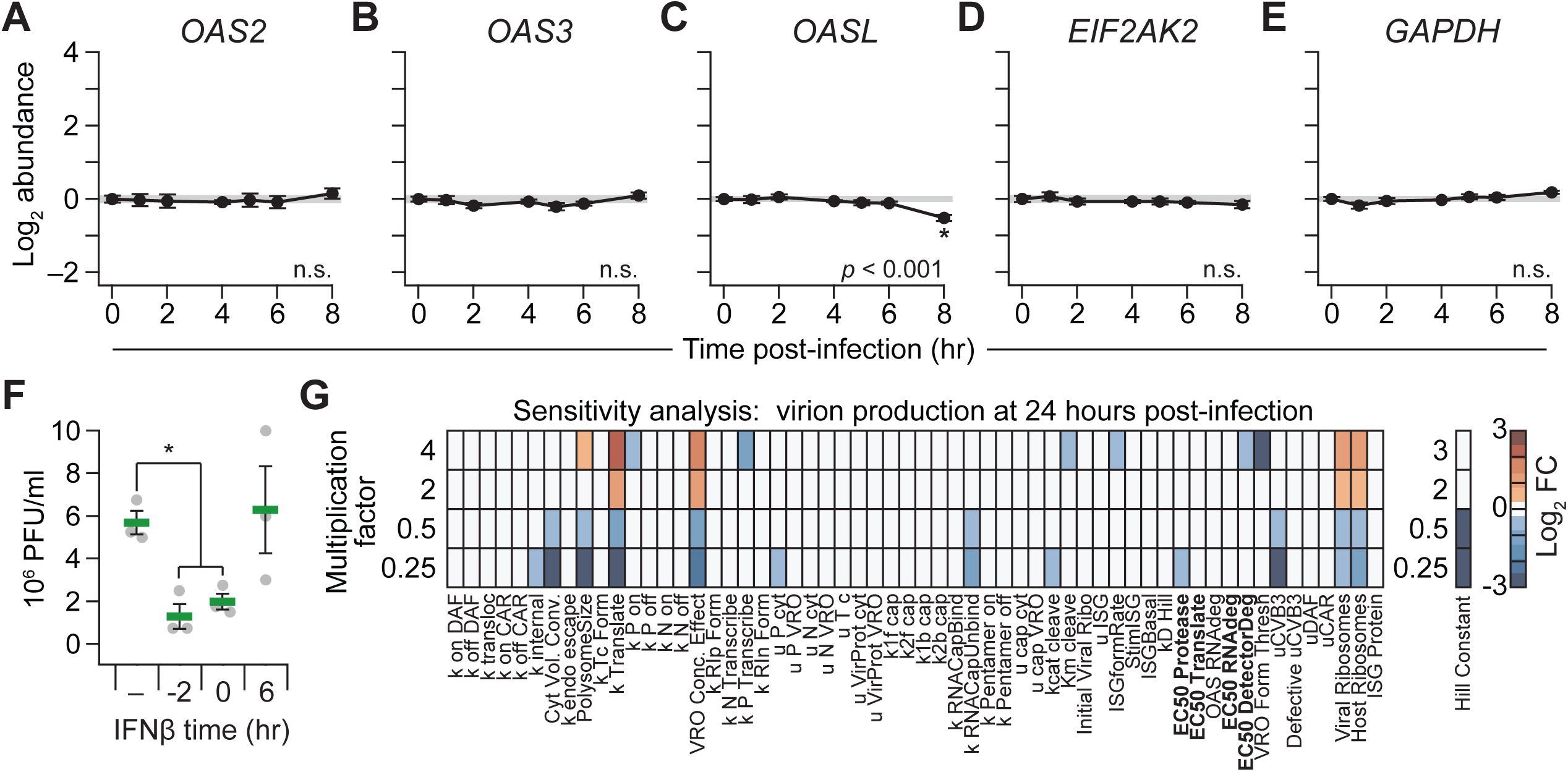
Related to Figure 5. Controls for interferon-stimulated genes and time-dependent antagonism of mature virion formation by supplemental interferon. (A–E) Total RNA was collected at the indicated times and measured for the indicated ISGs (*OAS2*, *OAS3*, *OASL*, *EIF2AK2*) or housekeeping gene (*GAPDH*) with *HINT1*, *PRDX6*, and *GUSB* used as loading controls. Data are shown as the geometric mean ± log-transformed standard error of *n* = 4 biological replicates. Time courses with significant alterations were assessed by one-way ANOVA with replication, and a single asterisk indicates *p* < 0.05 for individual time points compared to *t* = 0 hour after Tukey post-hoc correction. (F) Supplemental interferon blocks CVB3 progression during early infection. AC16-CAR cells were infected with CVB3 at MOI = 10 for 24 hours with 30 ng/ml IFNβ added at the indicated times relative to the start of CVB3 infection at *t* = 0, and the conditioned medium was assessed by plaque assay. Data are shown as the mean PFU/ml ± s.e.m. from *n* = 3 biological replicates. (G) Terminal virion production is not sensitive to the exact choice of half-maximal effective concentrations (EC50s, bolded) specifying feedback weights. The listed model parameters were individually varied by the indicated multiplication factor and the concentration of virions reported at 24 hours as the log_2_ fold change (FC) relative to the base parameter set. The Hill constant was varied multiplicatively below one and additively above one (where 2 = base Hill coefficient + 1).

**Figure S6.**
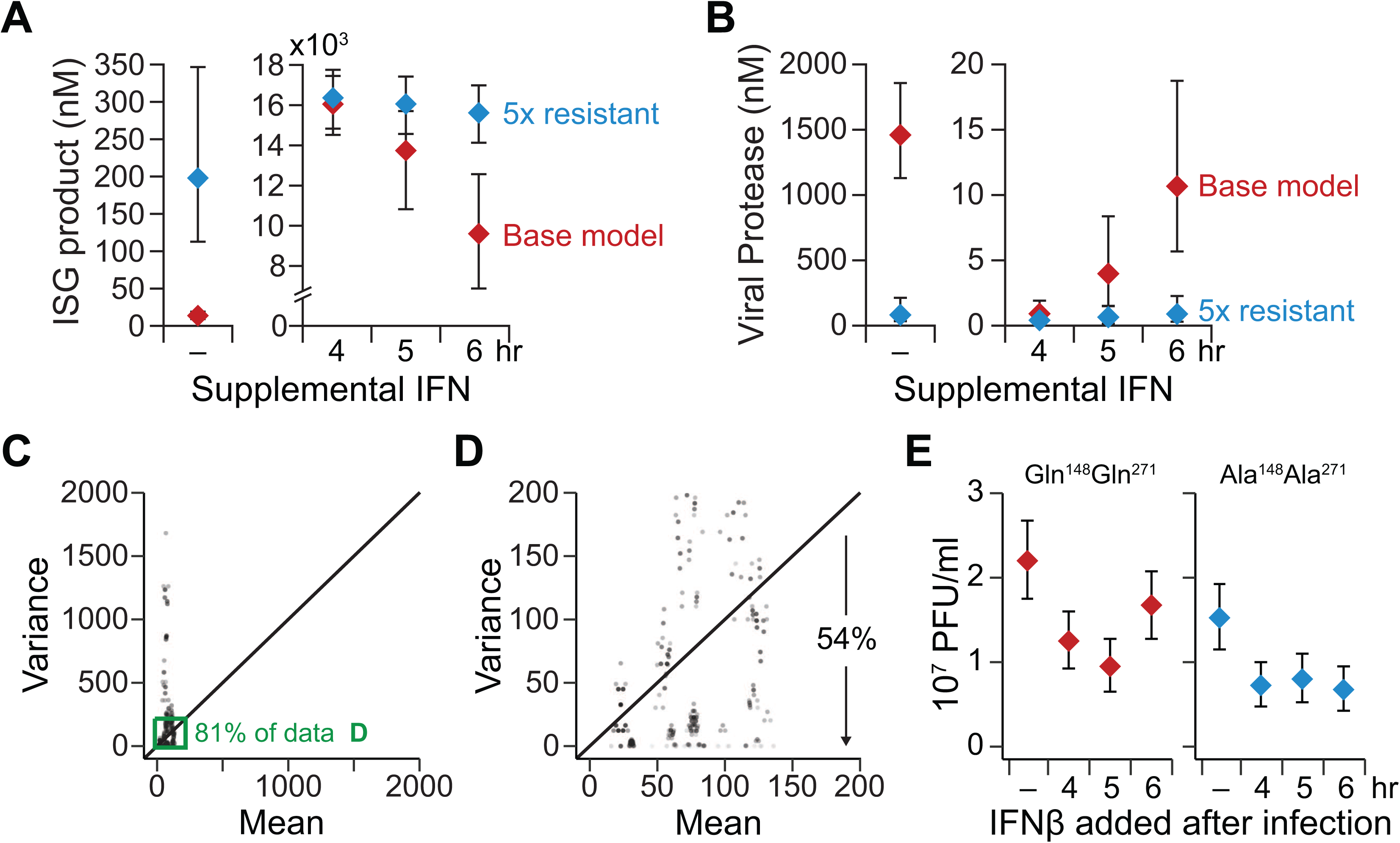
Related to Figure 6. Poisson noise of the plaque assay and generality of the timed-interferon effect to Ala^148^Ala^271^-mutant MAVS. (A and B) Predicted concentrations of ISG product and viral protease at 24 hours after infection with CVB3 and supplemental interferon (IFN) simulated at the indicated times. Predictions are shown as the median simulation ± 90% nonparametric confidence interval from *n* = 100 simulations of single-cell infections at 10 PFU with a parameter coefficient of variation of 5%. (C and D) Biological replicates of plaque assay are largely consistent with Poisson counting noise. After induction of LacZ or different MAVS alleles (Glu^93^Gln^148^Gln^271^, Glu^93^Gln^148^Ala^271^, or Gln^93^Gln^148^Gln^271^; STAR Methods), AC16-CAR cells were infected with CVB3 at MOI = 5 for 24 hours. Plaque assay data are from *n* = 4 biological replicates in one experiment (LacZ-induced cells), two experiments (Glu^93^Gln^148^Ala^271^ and Gln^93^Gln^148^Gln^271^ MAVS-induced cells), or three experiments (Glu^93^Gln^148^Gln^271^ MAVS-induced cells). Each experiment was bootstrapped 1000 times, yielding 8000 mean–variance pairs. The inset in (A) capturing 81% of the bootstrap replicates (green) is expanded in (B). (E) Ala^148^Ala^271^ MAVS qualitatively replicates Ala^271^ MAVS antiviral potency upon delayed addition of beta-interferon (IFNβ). After induction of Gln^148^Gln^271^ MAVS or Ala^148^Ala^271^ MAVS, AC16-CAR cells were infected with CVB3 at MOI = 5 for 24 hours, with 30 ng/ml IFNβ added at the indicated times after the start of CVB3 infection. Data are from one plaque assay at *n* = 4 different IFNβ conditions. Data are summarized as PFU/ml ± 95% Poisson confidence intervals based on the observation.

**Figure S7.**
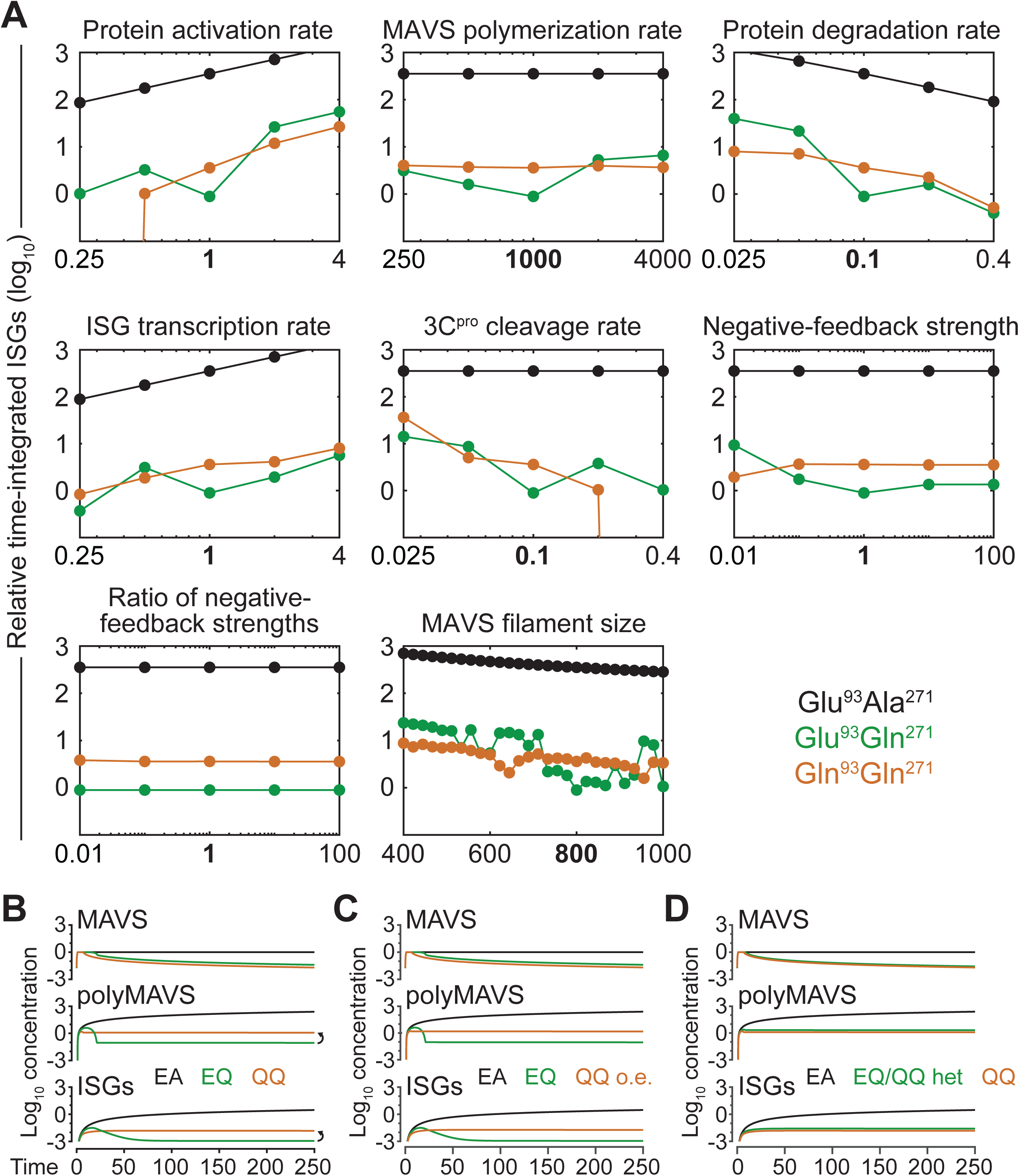
Related to Figure 7. Sensitivity analysis of the MAVS regulation model. (A) Individual model parameters were varied about their base value (bold) and simulations for the three genotypes repeated using integrated ISG abundance from 0–250 time units as the model output. (B–D) Effect of mixed MAVS genotypes. The original simulations from Figure 7F are reprinted in (B) for comparison to (C) a Glu^93^Gln^271^/Gln^93^Gln^271^ heterozygote overexpressing Gln^93^Gln^271^ ∼12- fold (QQ o.e., brown; Figure 6C) or (D) a Glu^93^Gln^271^/Gln^93^Gln^271^ heterozygote without overexpression (EQ/QQ het, green).

## KEY RESOURCES TABLE

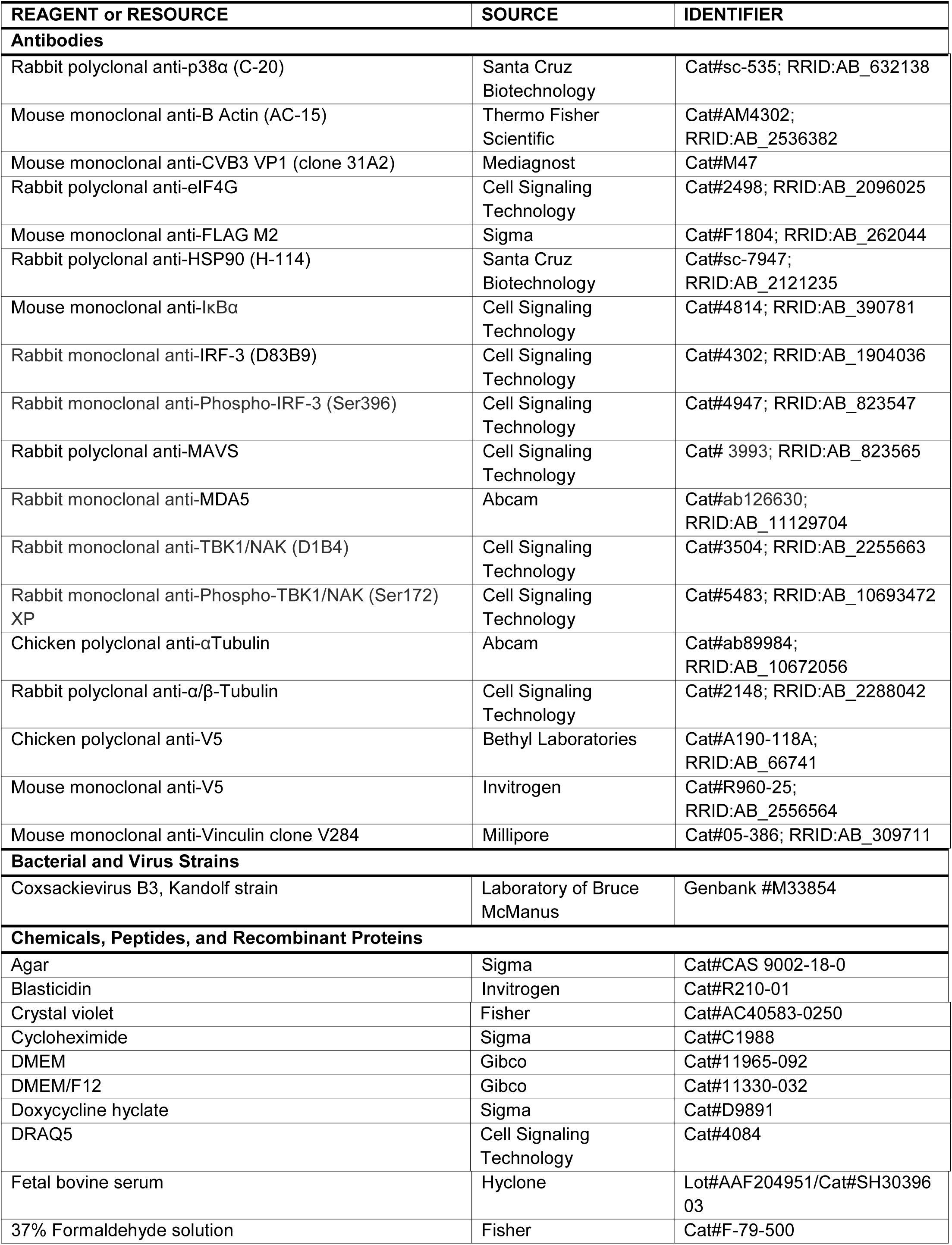
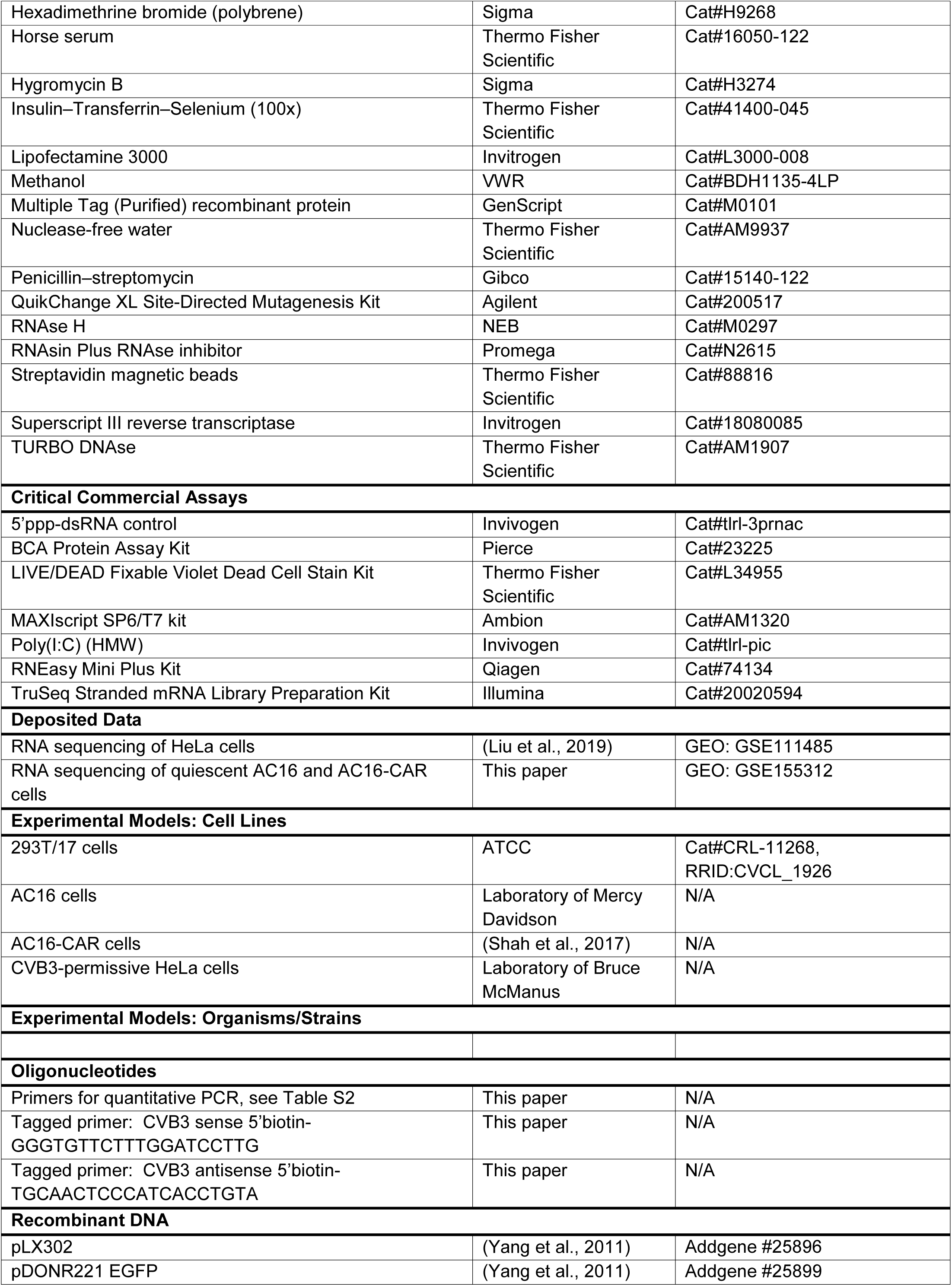
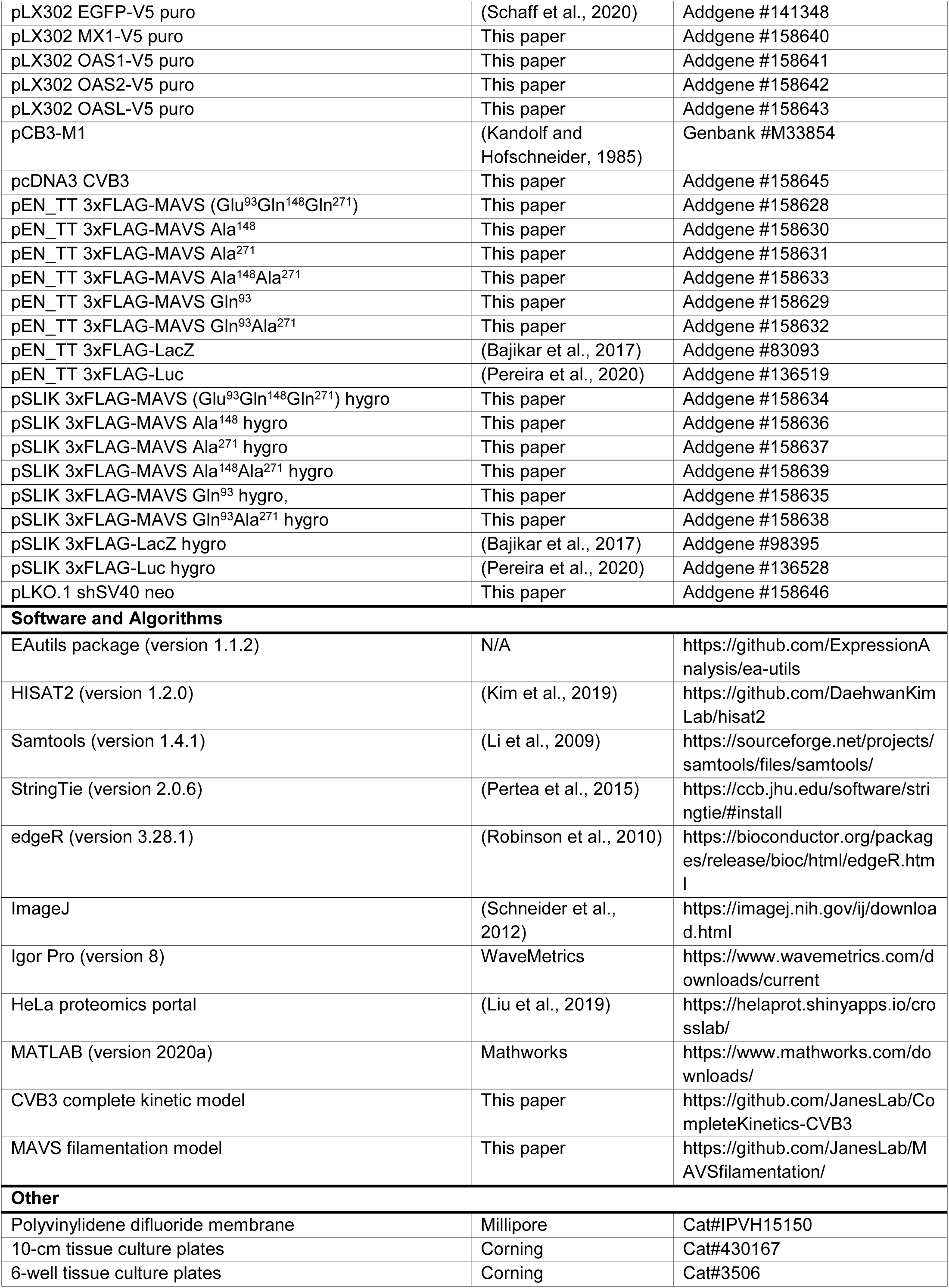

## Note S1. Related to Figure 7. Behavior of MAVS alleles in the MAVS filamentation model

The single-parameter sensitivity analysis indicated that time-integrated ISG profiles are robust to feedback inhibition of MAVS cleavage products on the uncleaved MAVS polymerization rate (Figure S7A). Feedback loops are thus omitted from this analysis of dynamical behavior, defined by a system of 4–6 differential equations depending on the MAVS allele:

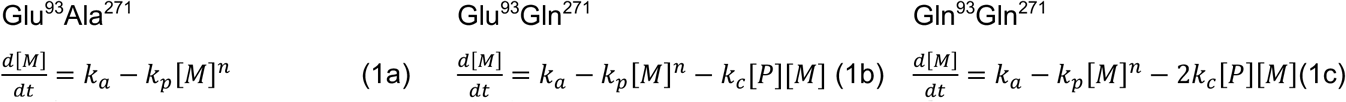

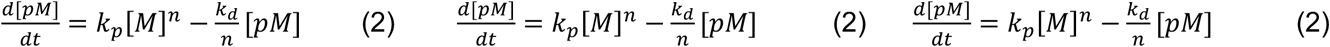

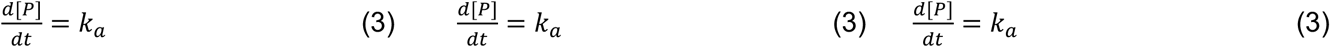

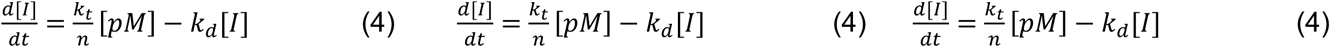

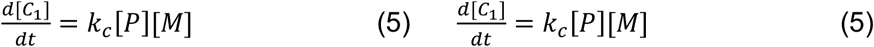

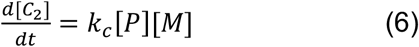

where *M* is uncleaved MAVS, *pM* is polymerized MAVS filaments, *P* is 3C^Pro^, *C_1_* is MAVS cleavage product 1, *C_2_* is MAVS cleavage product 2, *I* is ISGs, *k_a_* is a general activation rate, *k_p_* is the polymerization rate, *k_d_* is a general degradation rate, *k_t_* is the transcription rate, and *n* is number of MAVS monomers in a polyMAVS filament. For the same rate parameters, differences in the magnitude and duration of *I* depend entirely on the dynamics of *pM*, which depends on the dynamics of *M*. Further, the linear accumulation of *P* allows its analytical solution (*k_a_t*) to be substituted directly into the governing equation for *M*. Ignoring the bookkeeping equations of *C_1_* and *C_2_* reduces the system to two stable nonlinear, nonhomogeneous differential equations:

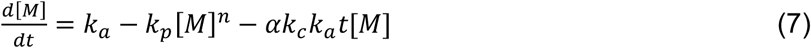

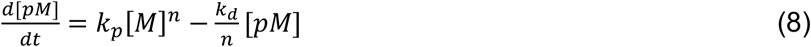

where *α* = 0 (Glu^93^Ala^271^), 1 (Glu^93^Gln^271^), or 2 (Gln^93^Gln^271^).

For *α* = 0, the long-term dynamics of the system can be solved analytically, with *M* rapidly reaching its pseudo steady-state value (*M_SS_*):

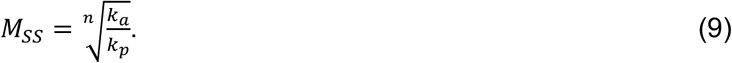

Substituting *M_SS_* into the governing equation for *pM* yields:

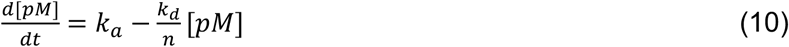

and the solution:

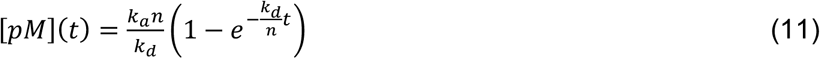

with *pM* approaching 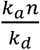 slowly with time, because *n* >> *k_d_*.

When *α* ≠ 0, an analytical solution does not exist. However, it is possible to identify critical transitions and limiting behavior of the dynamical system. For example, because *t* is always linearly increasing, one can define the time at which this initial *M_SS_* is no longer possible and *M* must decrease. Changing Equation (7) to an inequality:

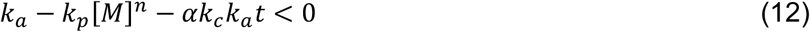

and solving for *t* yields a transition time for *M* (*t_M_*):

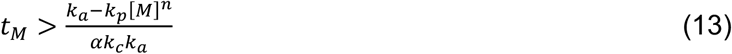

Thus, if the state of *M* is equal, the model predicts that Gln^93^Gln^271^ (*α* = 2) will transition twice as fast as Glu^93^Gln^271^ (*α* = 1; Figure SN1).

Immediately after *t_M_*, all terms in Equations (7) and (8) contribute to the dynamical-system behavior. However, the high multiplicity of MAVS polymerization implies that *M* will eventually decrease to a point that *k_p_[M]^n^* rapidly vanishes relative to the other terms in 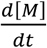:

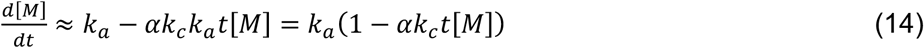

At this point, both *M* and *t* are large enough that we can further approximate Equation (14) as:

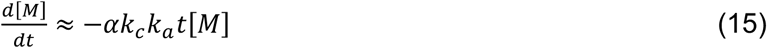

which is solvable analytically:

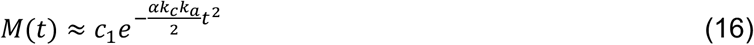

where *c_1_* is an unknown coefficient. Note from Equation (16) that the exponential decline occurs half as fast for Glu^93^Gln^271^ (*α* = 1) compared to Gln^93^Gln^271^ (*α* = 2). However, the approximate time point at which the *k_p_[M]^n^* term would vanish—shortly after *t_M_*—is twice as long for Glu^93^Gln^271^ according to Equation (13). Therefore, around the time when the system begins to deviate from steady state, the net decrease in *M* (and thus *pM*) occurs twice as fast for Glu^93^Gln^271^ as for Gln^93^Gln^271^ (Figure SN1).

Finally, at longer times, the rapid decline in *M* according to Equation (16) has progressed so much that it overwhelms the offsetting linear increase in *t* in Equation (14), and the simplification in Equation (15) is no longer valid. Here, the dynamics of *M* are governed by the linear, nonhomogeneous Equation (14).

**Figure SN1.**
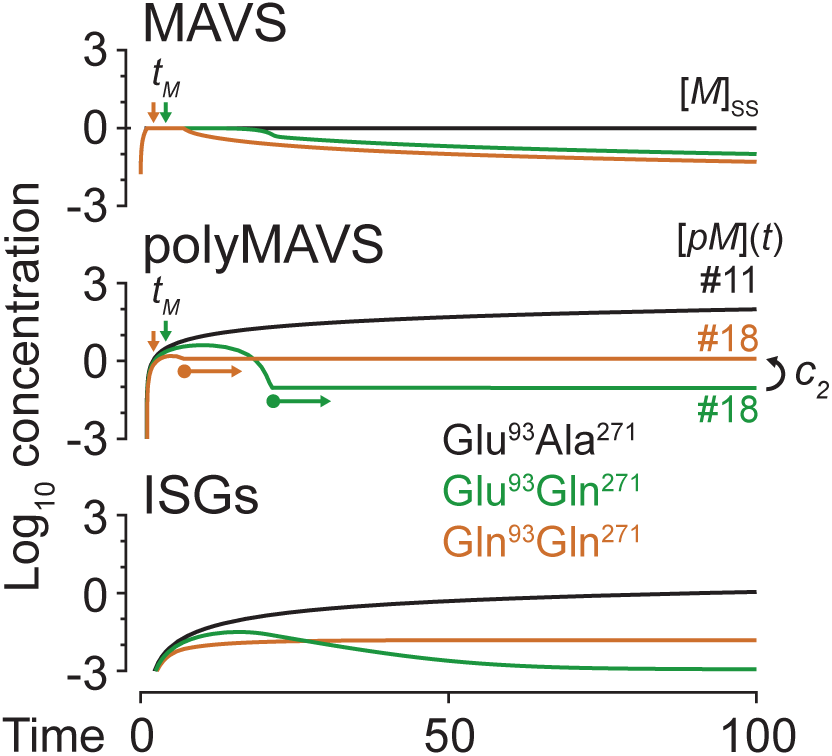
Annotated dynamics of the MAVS filamentation model. See text for description of variables and equations.

Numerical simulations suggest it is around this transition when *k_p_[M]^n^* vanishes relative to 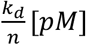 and the governing equation for *pM* [Equation (8)] can be approximated as:

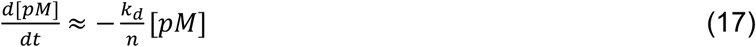

and solved to yield:

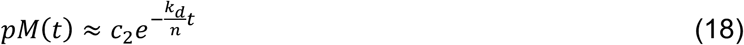

where *c_2_* is an unknown coefficient. In this regime, *pM* does not reach a formal steady state but is operationally stable at *c_2_* because *n* >> *k_d_* (Figure SN1). For Glu^93^Gln^271^, the delayed transition from steady state [Equation (13)] causes a faster rate of decline in *M* [Equation (16)], leading to a deeper plunge in *pM* before the transition to an operationally stable concentration [Equation (18)].

**Table S2.**
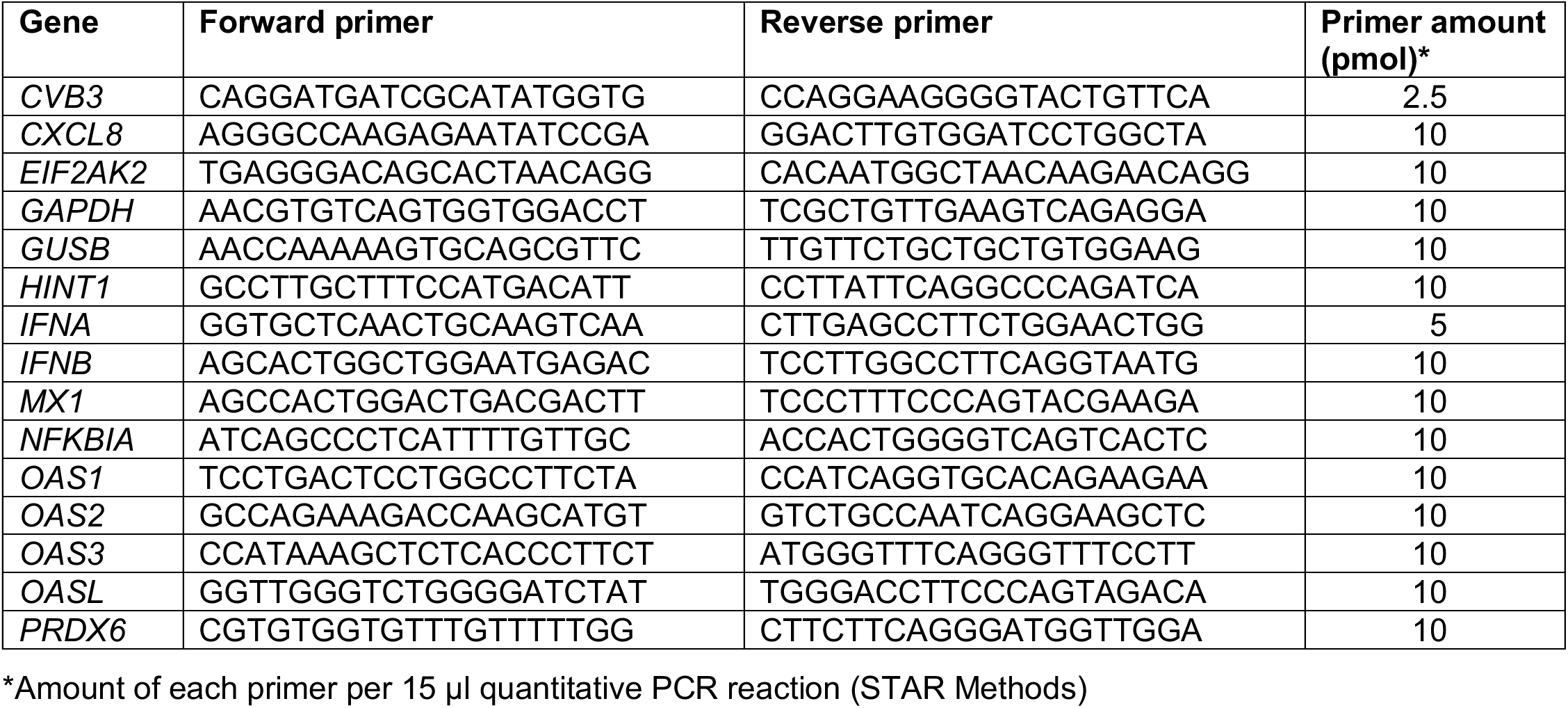
Related to STAR Methods. Quantitative PCR primers used in this study.

